# Thyroid hormone receptor beta mutations alter photoreceptor development and function in *Danio rerio* (zebrafish)

**DOI:** 10.1101/800821

**Authors:** Ciana Deveau, Xiaodong Jiao, Sachihiro Suzuki, Asha Krishnakumar, Takeshi Yoshimatsu, J Fielding Hejtmancik, Ralph F. Nelson

## Abstract

We investigate a splice variant of *thrb* isolated in the retina, *trβ2*, identifying functional changes in larval and adult mutant zebrafish lacking trβ2. We constructed two CRISPR mutant zebrafish with mutations located in the N-terminus region. The first is a *6BP*+*1* insertion deletion frameshift resulting in a truncated protein. The second is a *3BP* in frame deletion with intact binding domains. ERG recordings showed that the *6BP*+*1* mutants did not respond to red wavelengths of light while the *3BP* mutants did respond. *6BP*+*1* mutants lacked optomotor and optokinetic responses to red/black and green/black contrasts. Adult *6BP*+*1* mutants exhibit a loss of red-cone contribution to the ERG, and an increase in green and UV contributions. Anatomical markers show loss of red-cones in the *6BP*+*1* mutant but increase in blue, green, and UV cone density. Our results confirm *trβ2*’s role in retinal cone development.

**Author Summary:** There are four cone photoreceptors responsible for color vision in zebrafish: red, green, blue, and UV. The thyroid hormone receptor *trβ2* is localized in the vertebrate retina. We know that it is necessary for the development of long-wavelength-sensitive cones (red), but here we investigate the functional alterations that accompany a loss of *trβ2*. Our work contributes to the ongoing investigations of retinal development and the involvement of thyroid hormone receptors. Confirming previous morphological findings, we see that the fish become red colorblind when *trβ2* is knocked out, but the contributions of the other three cone types shift in response. Our work highlights the plasticity of the retinal circuit as we see changes in opsin peaks and cone sensitivity, increases in contributions of UV cones, and an attempt at a mosaic pattern in the adult retina all in the absence of *trβ2* and red cones. We now have an increased understanding of mechanisms underlying retinal development

## Introduction

To achieve vertebrate color vision, cones with different wavelength sensitivities develop from retinal progenitor cells. Cone signals stimulated by their respective wavelengths of light merge post-synaptically to create the visual perception of color in the brain. Proper differentiation of these cones, and subsequent proper color perception, is reliant on specific transcription factors. The vertebrate family of long-wavelength-sensing (LWS) cones are linked to thyroid hormone receptors. Differentiation of LWS cones in rodents requires an isoform of thyroid hormone receptor beta (*thrb*) called *trβ2* [1], one of two thyroid hormone receptors in a family of nuclear receptors [2]. Human heterozygous *thrb* mutants manifest a metabolic syndrome, resistance to thyroid hormone (RTHβ). In RTHβ there are color vision deficits [3]. In one reported human *thrb* homozygous mutant, both LWS and middle-wavelength-sensitive (MWS) cone functions were severely depressed, while short-wavelength-sensitive (SWS) cone function was enhanced [4]. Thyroid receptors (TRs) are ligand-dependent transcription factors that contain an N-terminus, a DNA binding domain, and a ligand (T3) binding domain, conserved across multiple vertebrates including chicken, mouse, human, and zebrafish [5, 6, 7, 8]. *Thrb* is broken down into multiple isoforms through alternative splicing, which are active in the vertebrate retina, pituitary gland, and inner ear [9]. The *trβ2* isoform is localized in the retina [9, 10].

Zebrafish (*Danio rerio*) is an important color vision model. It has four opsin types, LWS, RH-2, SWS2, and SWS1 [11]. Zebrafish has two opsins derived from the LWS evolutionary tree by gene duplication (*LWS1* & *LWS2*), and one from the SWS or UV-opsin evolutionary tree (*SWS1*). Humans also express two homologous LWS opsins (*LWS* & *MWS*), and one SWS opsin (*SWS*). In both species the two LWS opsins are in a tandem gene array governed by a common promoter region [12]. Each cone type expresses only one class of opsin in zebrafish [11]. *Trβ2* was previously shown in zebrafish to be essential for the development of red-cones and sufficient for inducing L-opsin expression with transgenic overexpression leading to an increase in LWS (red) cones and a decrease in SWS1 (UV) cones [10]. Our aim is to investigate germline alterations of *trβ2* expression. Although a morpholino knockdown of *trβ2* in 5 day larvae leads to a decrease of red-cones and corresponding increase of UV cones [13, 10], it is unknown how that disrupts color vision functionally in larvae, and adult mutants are unexplored. This paper is the first to relate retinal physiology with behavior and study the spectral tuning of cones with respect to the *trβ2* mutation.

To answer those questions, we established two *trβ2* mutant lines using the CRISPR/Cas9 system. The first is a *c.184_188delTATGGinsGTTCCC* (*6BP*+*1*) frameshift indel and the second is a *c.184_186delTAT* (p.Tyr61del) (*3BP*) a single in-frame codon deletion, both located in the first exon of the *trβ2* isoform, which is unique to this isoform and thus would be expected to affect only retinal development. The *6BP*+*1* frameshift mutation creates an early stop codon, eliminating the DNA binding site and ligand binding site. The *3BP* mutation deletes Tyr61, an N-terminus amino acid that is highly conserved among vertebrates, but leaves the DNA binding site and ligand binding site intact. The *6BP*+*1* mutation resulted in the anticipated LWS-cone loss, but the *3BP* mutation revealed unexpected results. Here we characterize the physiological differences through spectral ERGs of the PIII cone signal in zebrafish, changes in color perception through optomotor and optokinetic behavioral testing, and antibody staining for key cone markers to provide a broad picture of the functional, behavioral, and molecular alterations. We found that the changes in the makeup of the cone layer were reflected in both the ERG responses and behavior, supplementing our knowledge on the role of *trβ2* in the retina.

## Results

### CRISPR mutations are located in the N-terminus of trβ2

The two splice variants of the *thrb* gene, *trβ1* and *trβ2* differ only in the first exon [14]. To target *trβ2*, we designed gRNA targeting exon1 of *trβ2* and injected into one-cell-stage eggs of tg(*trβ2*:tdTomato). We isolated two mutations of *trβ2*. Mutations were found at the CCC (proline) TAT (tyrosine) juncture. (Fig. 1A-C). The mutation in Figure 1B deletes a TATGG and replaces it with a GTTCCC, a frameshift indel mutation leading to a stop codon in the first exon. We refer to this as the *6BP*+*1* mutation. The mutation in Figure 1C deletes a single codon, TAT (tyrosine), from the *trβ2* sequence. We refer to this as the *3BP* mutation. The mutations occurred in the first exon of *trβ2*, in a region specifically involving the Tyr61 in the N-terminus, at the focus of the CRISPR/Cas9 targeting sequence. The *6BP*+*1* mutation eliminated both the DNA binding site (DBD) and thyroxin ligand binding site (LBD). The *3BP* mutation left both those regions intact but altered the protein binding site (PBD) at a highly conserved amino acid, possibly involved in transcription factor activation [5]. The germ-line genetic loss of functional *trβ2* is necessary to study phenotypes beyond the larval stage. Morpholinos are only useful for experiments up to ∼5dpf because the gene suppression is transient so we developed the *trβ2* mutant line.

**Fig 1:**
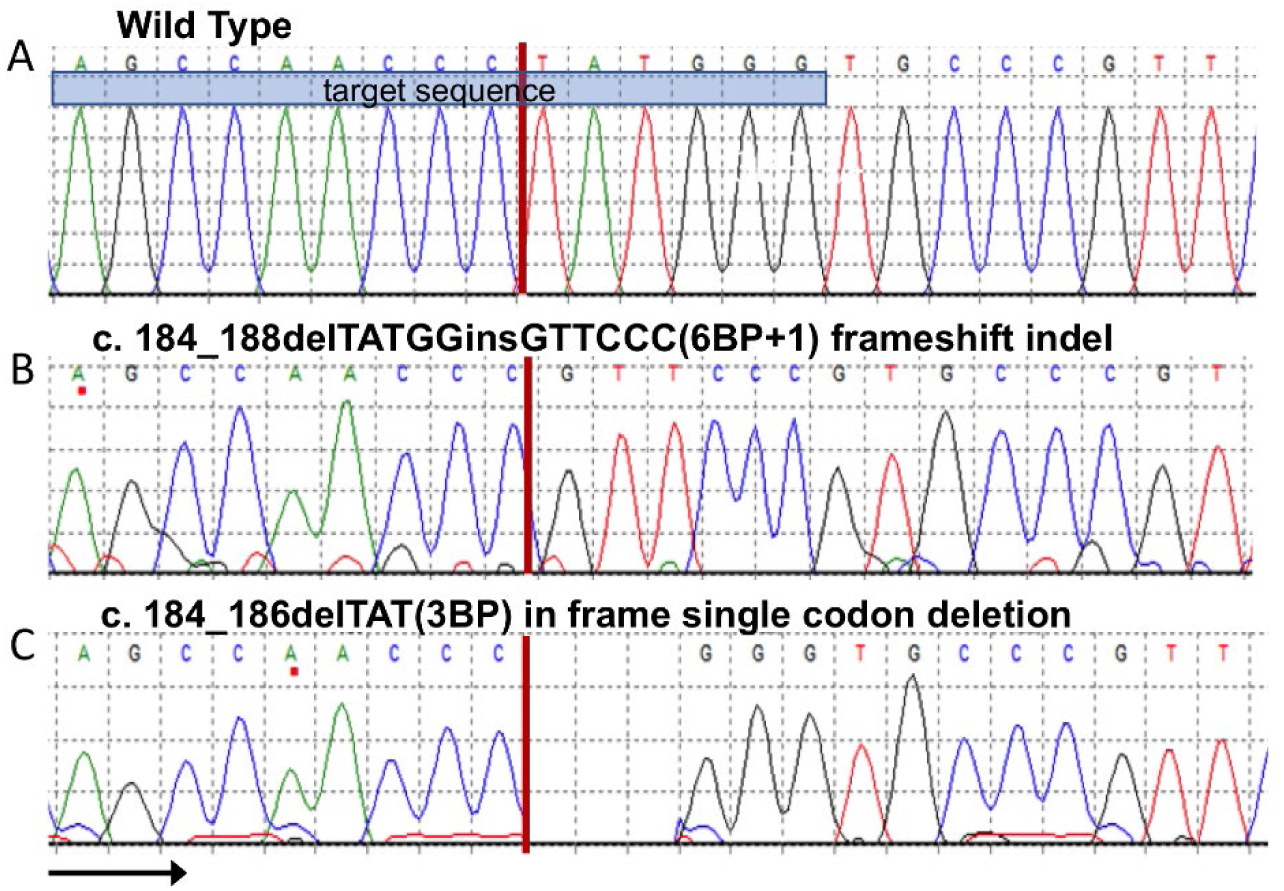
Two CRISPR mutant lines. (A-C) Both mutations targeted the TAT codon resulting in a frameshift indel (6BP+1) and an in-frame codon deletion (3BP). DNA binding domain (DBD), ligand binding domain (LBD).

### Gene reporter activity for trβ2 in trβ2 mutants and heterozygous mutants

The *trβ2* gene is thought to promote its own transcription. Therefore, the absence of competent *trβ2* protein product is expected to diminish promoter activity for the *trβ2* gene. To test this we crossed *6BP*+*1* fish carrying the reporter transgenes *trβ2:*tdTomato with *6BP*+*1* heterozygous fish. Similarly, we in-crossed *3BP* heterozygous fish carrying *trβ2:*mYFP [10]. Larvae were raised in PTU to block pigment epithelial melanin formation so that the fluorescent products of the reporter transgenes could be visualized within the larval eyes using live confocal imaging (Fig. 2). The transgenes produce fluorescent red-cone layers in the *6BP*+*1* WT (+/+, Fig. 2A) and the *6BP*+*1* heterozygous mutant (+/-, Fig. 2B) but not the *6BP*+*1* mutant (-/-, Fig. 2C). For fluorescence phenotyping of the *6BP*+*1* strain all 11 cone-bright phenotype images were either a heterozygous (+/-) mutant or wild type (+/+) genotype, while mutant (-/-) genotypes failed to fluoresce. This indicates that either a full (+/+) or half (+/-) dose of native *trβ2* is enough to activate the *trβ2* promoter, but the *6BP*+*1 trβ2* protein, truncated in the first exon, is not. The result further affirms that functional *trβ2* protein is necessary for transcription of the *trβ2* gene.

**Fig 2:**
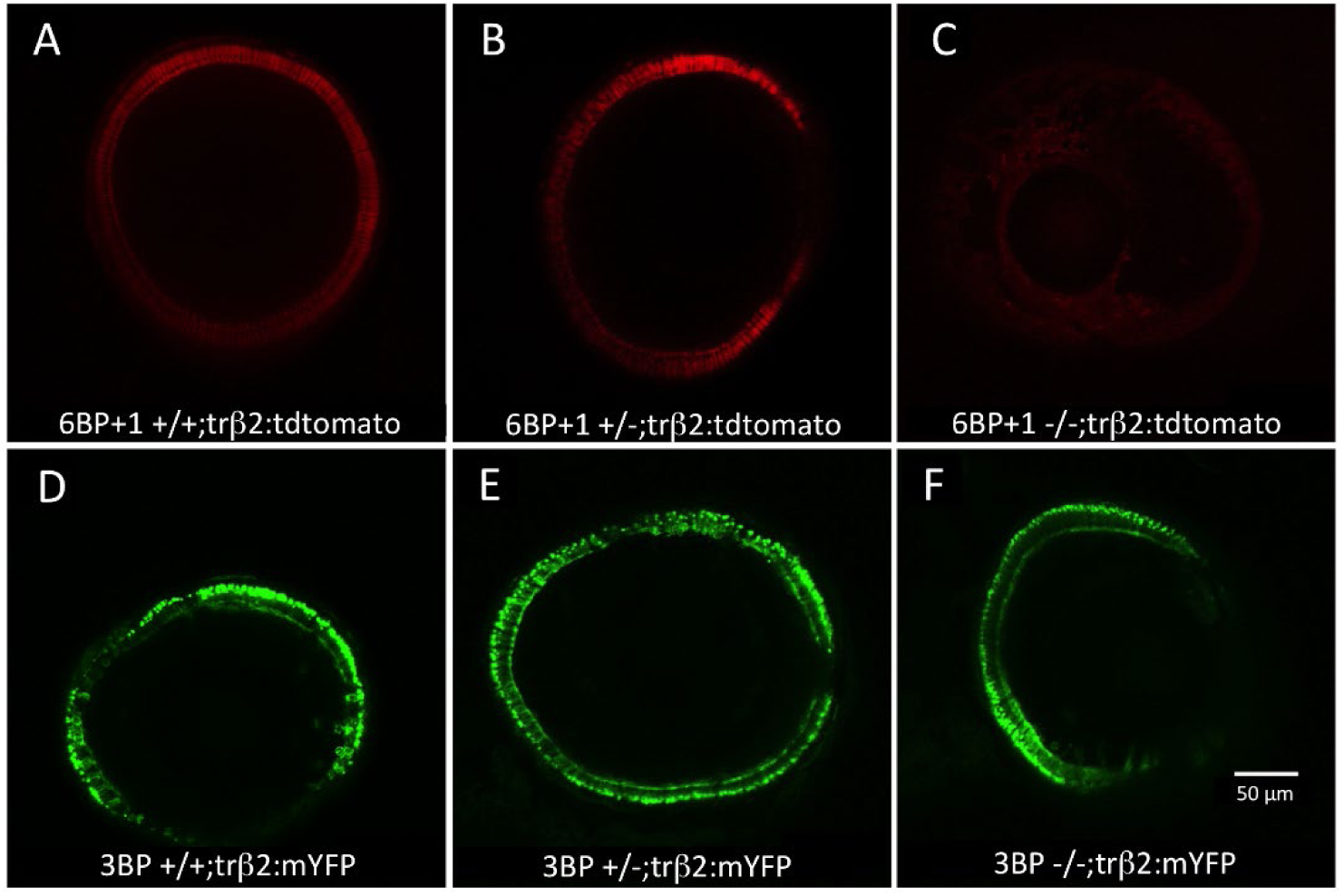
Live confocal imaging of larval eyes at 4- and 5-days post fertilization (dpf). (A) Wild type larval retina has a clear red-cone structure fluorescing by tdTomato with a *trβ2* promoter. (B) A 6BP+1 heterozygote maintains *trβ2* fluorescence, but the 6BP+1 mutant (C) has no *trβ2* fluorescence. The 3BP wild type (D), heterozygote (E), and mutant (F) all have *trβ2* fluorescence shown through mYFP with a *trβ2* promoter.

In confocal imaging, there is no clear difference between the larval cone layer fluorescence in the *3BP* mutant (-/-, Fig. 2F) versus the het mutant (+/-, Fig. 2E) and the wild type (+/+, Fig. 2D). Fluorescent reporter activity based in the *trβ2* promoter is evident in all three genetic lineages of the *3BP* heterozygous in-cross. Despite the absence of the highly conserved tyrosine, the *3BP trβ2* protein activates the native *trβ2* promoter.

### Immunohistochemistry on larval and adult retinas show a complete loss of red cones and opsin in the 6BP+1 mutants

Larval retinas fixed at 7 dpf were stained for red opsin on a DAPI background. The *6BP*+*1* mutant did not show red opsin antibody fluorescence or tdTomato transgene fluorescence (Fig. 3A). In contrast, the *6BP*+*1* heterozygote expressed red opsin antibody fluorescence co-localized with *trβ2*:tdTomato expression (Fig. 3B). This mutant knockout result is consistent with previous morpholino knockdowns of *trβ2* in larvae [10, 13].

**Fig 3:**
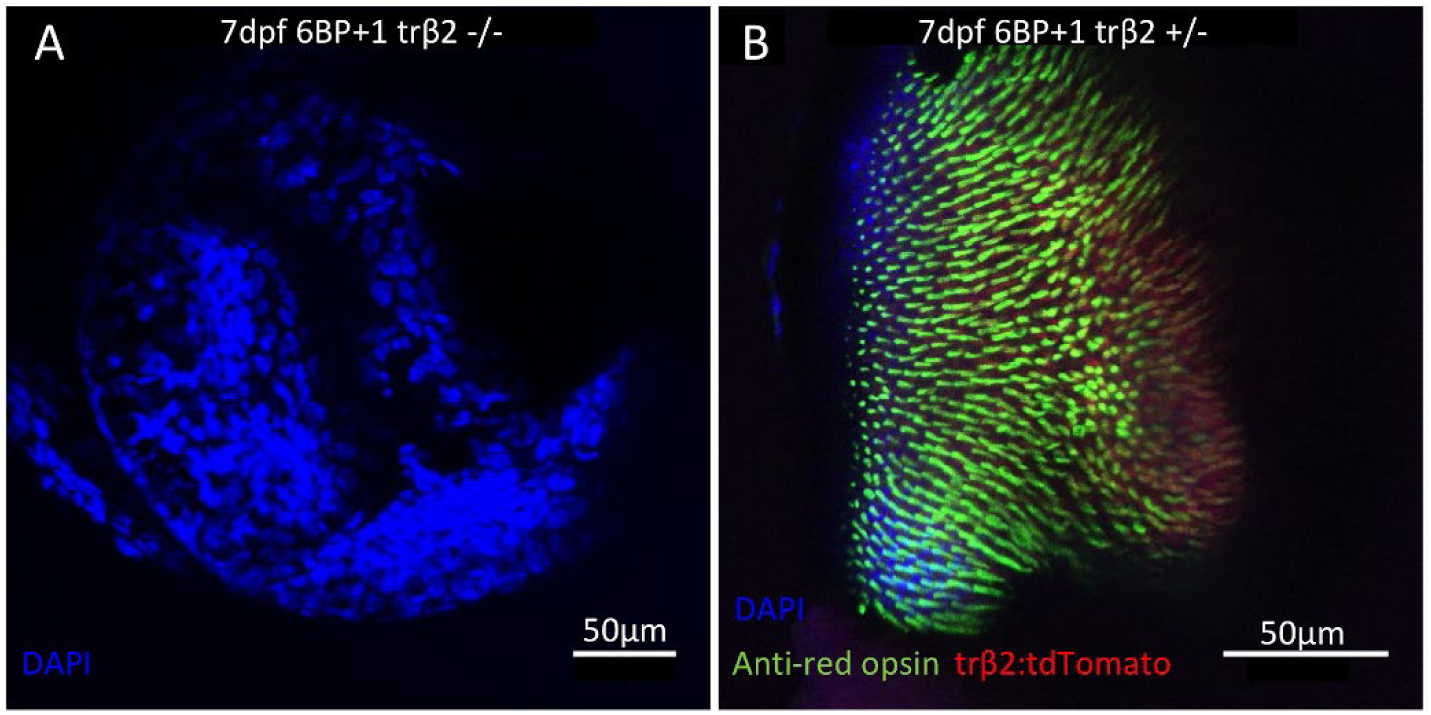
Antibody stains of 7 dpf larval retinas. (A) Mutant retina did not have red opsin or *trβ2* fluorescence. (B) Heterozygote retina showed the red opsin staining colocalized with tdTomato.

This loss of red cones extends into adulthood. Retinas of adult fish between 10- and 12-months post fertilization were stained with the arrestin (arr3) antibody, zpr-1, marking both red principal and green accessory members of double cones and also a red opsin antibody (1D4). The *6BP*+*1* mutant had a complete loss of both red markers (Fig. 4A, D, G), while the heterozygote and wild type retinas have overlapping and distinct double-cone and red opsin fluorescence (Fig. 4B, C, E, F, H, I). A *trβ2* chimeric transgenic shows a distinct loss of red cones in the *trβ2-/-* region (right) of the retina (Fig. 5D). Green and blue cone patterning is altered in the mutant region as there appears to be close proximity between these two cones (Fig. 5C, 6B), while in the wild type, blue cones only associate with red cones [15]. The mosaic ratio of the *trβ2*+*/*+ region is consistent with previous findings of 2:2:1:1 (R:G:B:UV) [15]. While the ratio between the green, blue, and UV cones remain the same in the mutant (2:1:1), there is a significant increase in the number of each of those cone types (Fig. 6). Cell counts were taken from five 45µm by 45µm sections in both regions of the chimeric retina and averaged. Green cones increased from 72.60±3.53 to 91.20±2.20 (t-test, *df* =8, P=0.0021), blue cones increased from 35.80±1.32 to 49.20±0.49 (t-test, *df*=8, P<0.0001), and UV cones increased from 35.60±2.21 to 48.80±1.02 (t-test, *df*=8, P=0.0006). There is still a blue/UV cone row and a green cone row observed in the mutant, though it is not as clearly distinguished as it is with the red cones (Fig. 6B).

**Fig 4:**
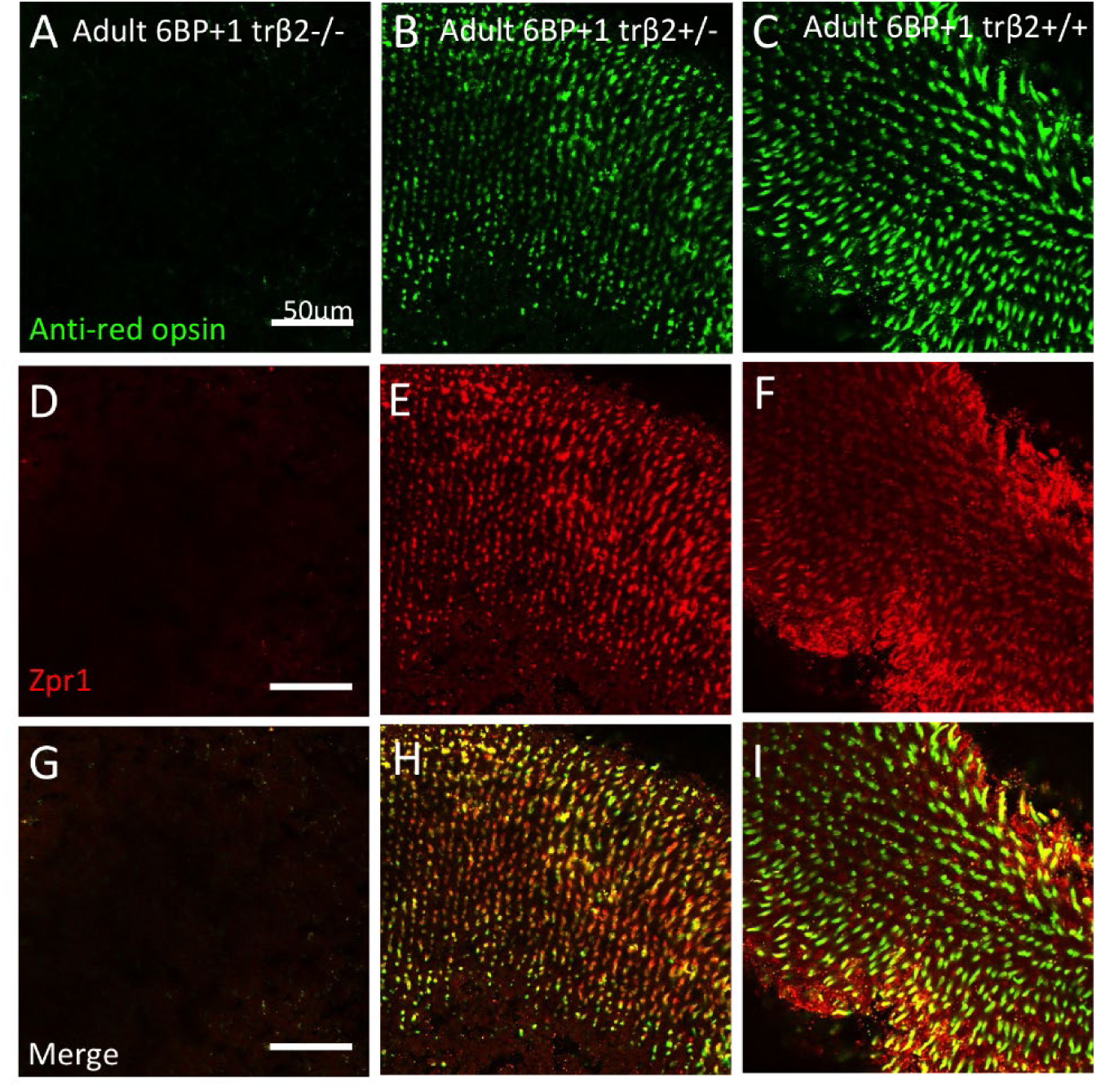
Anti-red opsin and zpr1 antibody staining of 6BP+1 adult. (A, D, G) Mutant adult retina did not have red opsin or red-green double cone fluorescence. (B, E, H) Heterozygote adult retina showed bright staining for red opsin and red-green double cones, as seen in the wild type (C, F, I). Scale is the same for each image.

**Fig 5:**
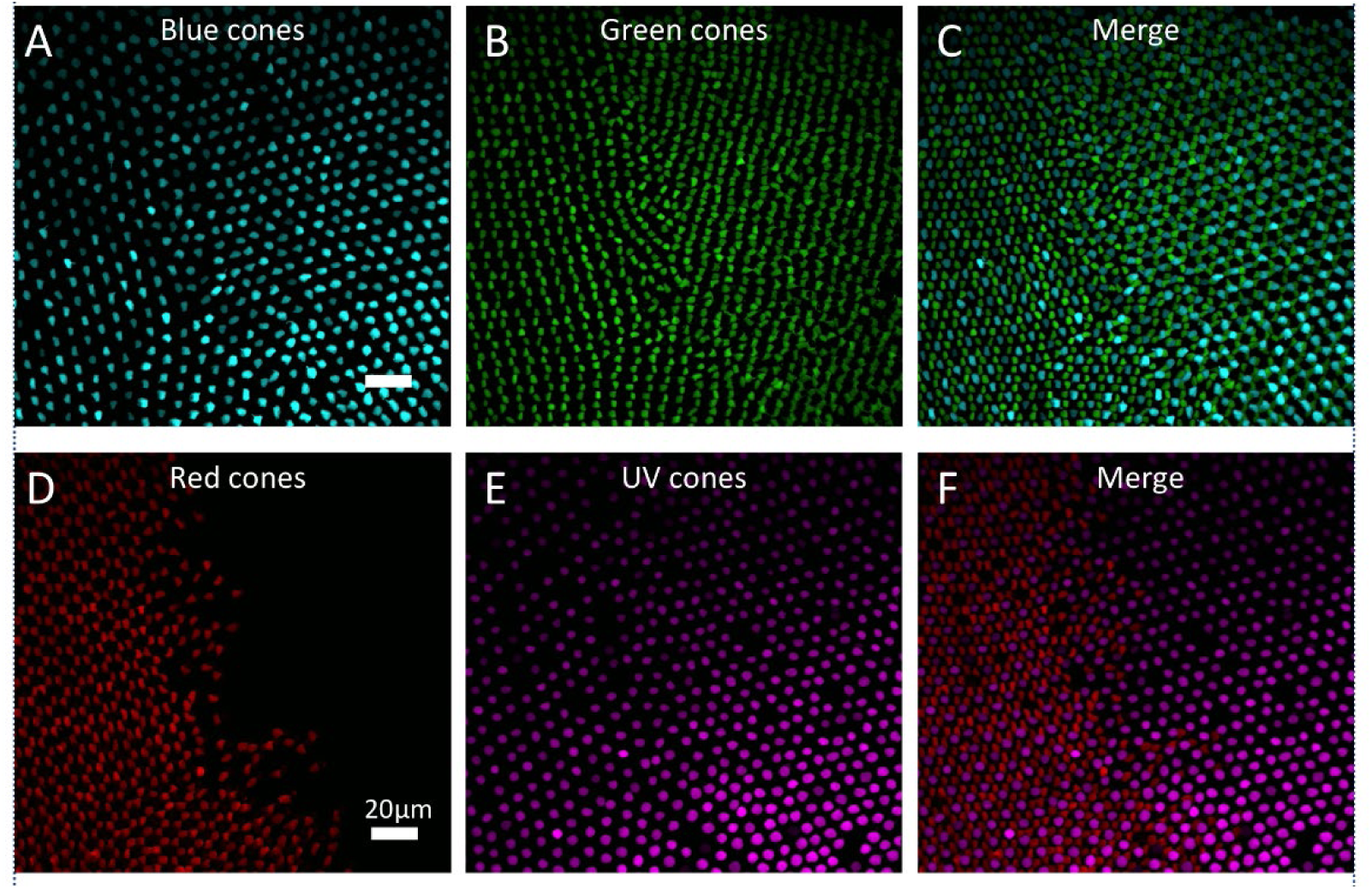
Cone photoreceptor spatial arrangements of 21-day *trβ2* chimera. Left side of each image is either wild type or *trβ2*+*/-*. Right side of each image is *trβ2-/-*. Images show cone patterning separately (A, B, D, E) and merged (C, F). Scale is the same for each image.

**Fig 6:**
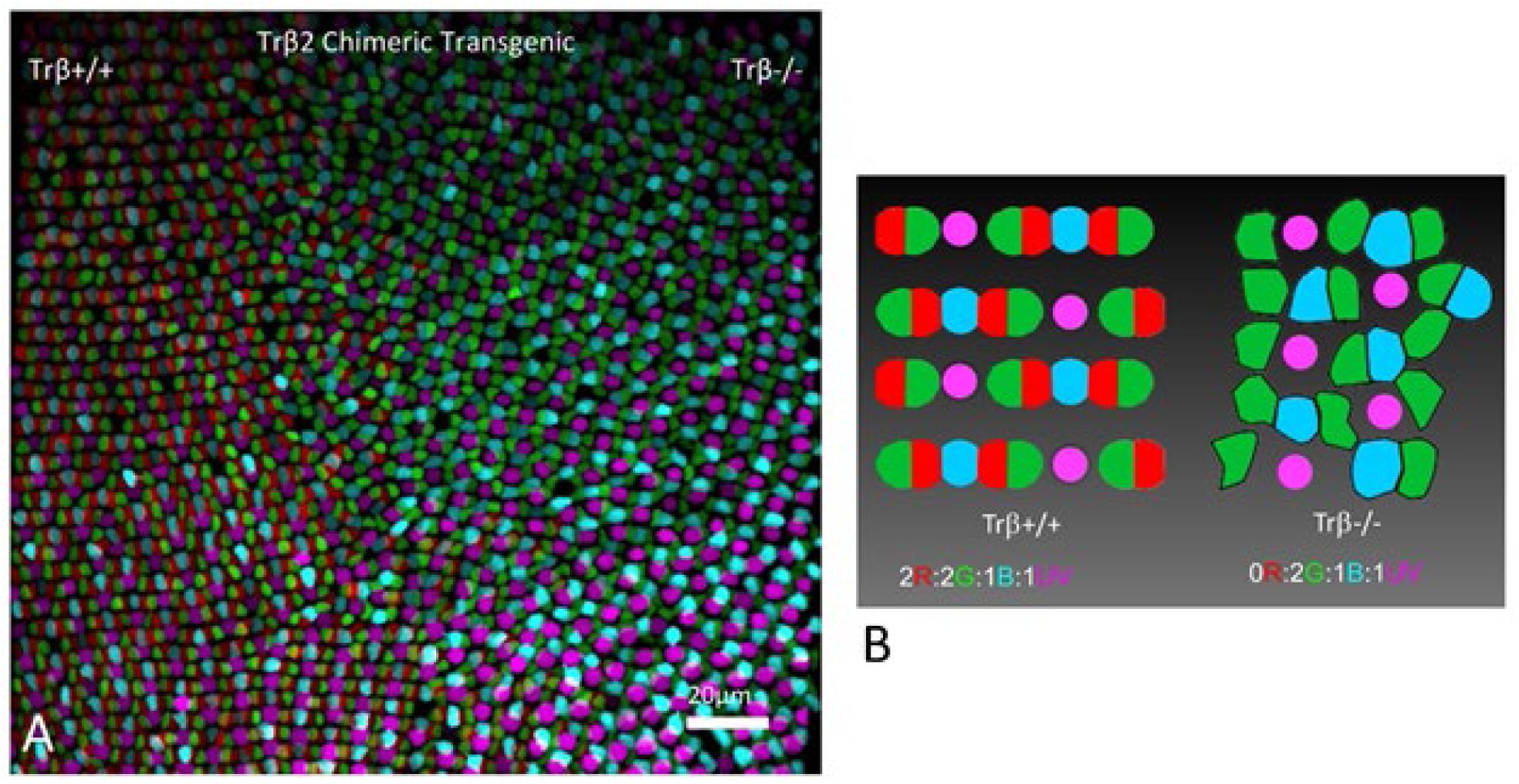
Mosaic analysis of cone photoreceptor spatial arrangement with CRISPR/cas9-mediated genome editing of *trβ2*. (A) In this 21 dpf chimeric the left side of the retina is the wild type or *trβ2*+*/-*, while the right side is *trβ2-/-*. (B) Cone mosaic patterns in wild type/*trβ2*+*/-* (left) or *trβ2-/-* (right). Red cones (red), green cones (green), blue cones (blue), UV cones (magenta)

### Zebrafish larvae with the 6BP+1 trβ2 mutation have a significant loss of response to red light and altered fractional cone contributions

We blocked signals downstream of the cones to extract the larval cone PIII waveforms. These signals in response to spectral stimulation for a *trβ2 6BP*+*1* mutant eye (-/-) and a *trβ2 6BP*+*1* heterozygous mutant eye (+/-) are shown in Figure 7. Each set of 7 nested traces is an irradiance series, with 0.5 log unit increments in stimulus brightness, and with wavelengths as shown in each set delivered consecutively in left to right time order. The cone PIII waveform consists of vitreal hyperpolarization during the light stimulus (rectangular trace) with a vitreal depolarizing rebound peak following stimulus offset. No b-waves appear in these records, as the activity of synapses on second order retinal neurons is blocked by 20 mM aspartate.

**Fig 7:**
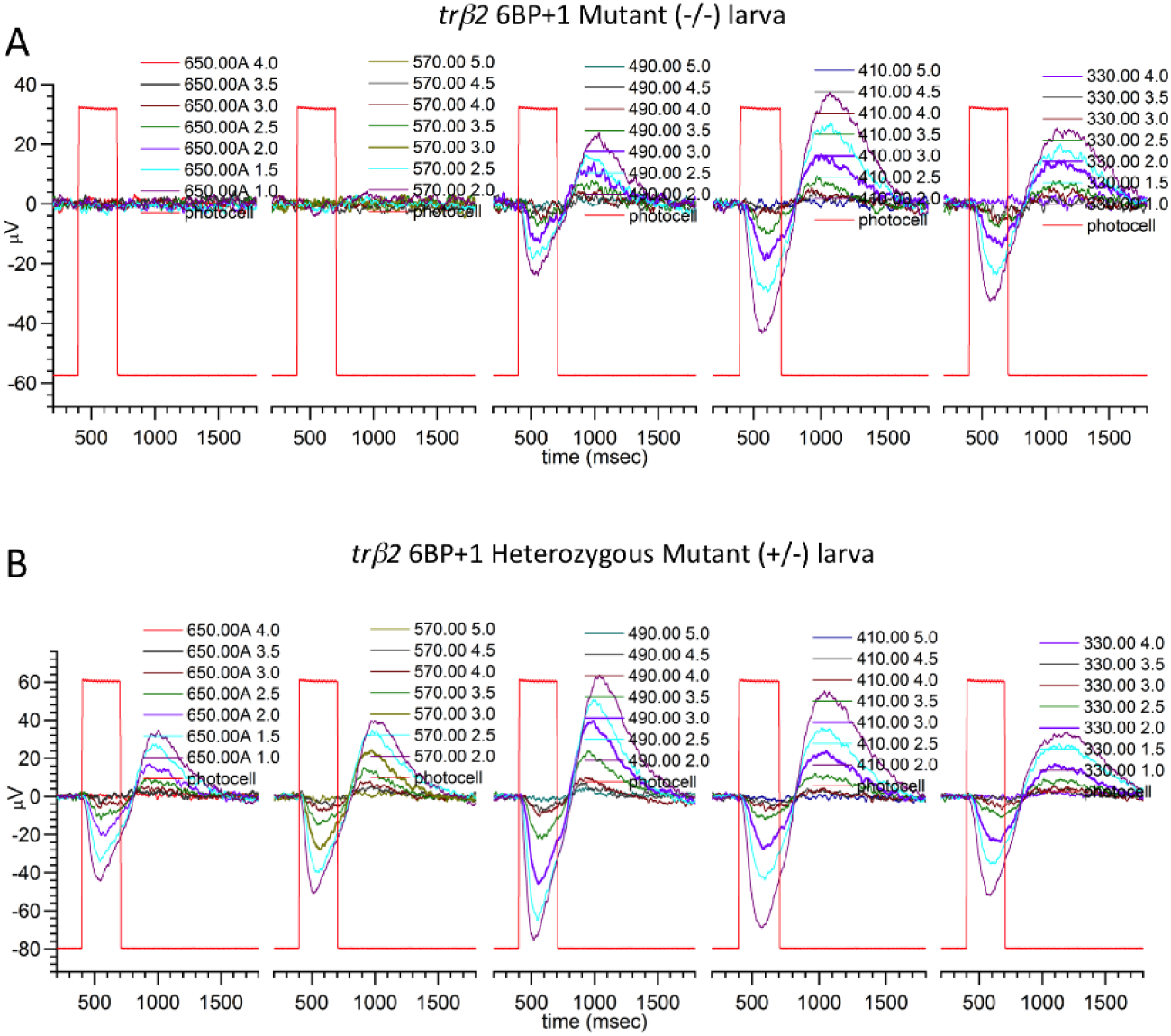
*6BP*+*1* larval cone PIII ERG spectral response traces. (A) The *6BP*+*1* mutant does not respond to 650 nm and 570 nm while maintaining a response to shorter wavelengths. (B) The heterozygote shows sustained responses across all wavelengths (650 - 330 nm). Larvae are 5-7 dpf. Cone PIII responses isolated with 20mM Na Aspartate

In the *trβ2 6BP*+*1* -/- eye (Fig. 7A), PIII ERG responses are only seen for wavelengths shorter than 570 nm. The *trβ2 6BP*+*1* +/- eye responds at all wavelengths (Fig. 7B). This suggests the absence of red (LWS) cone signals in the mutant, but not the heterozygous mutant eye. In a larval *trβ2 3BP* -/- eye (Fig. 8A), light responses appear at all stimulus wavelengths, as they do in a wild type (WT) eye (Fig. 8B).

**Fig 8:**
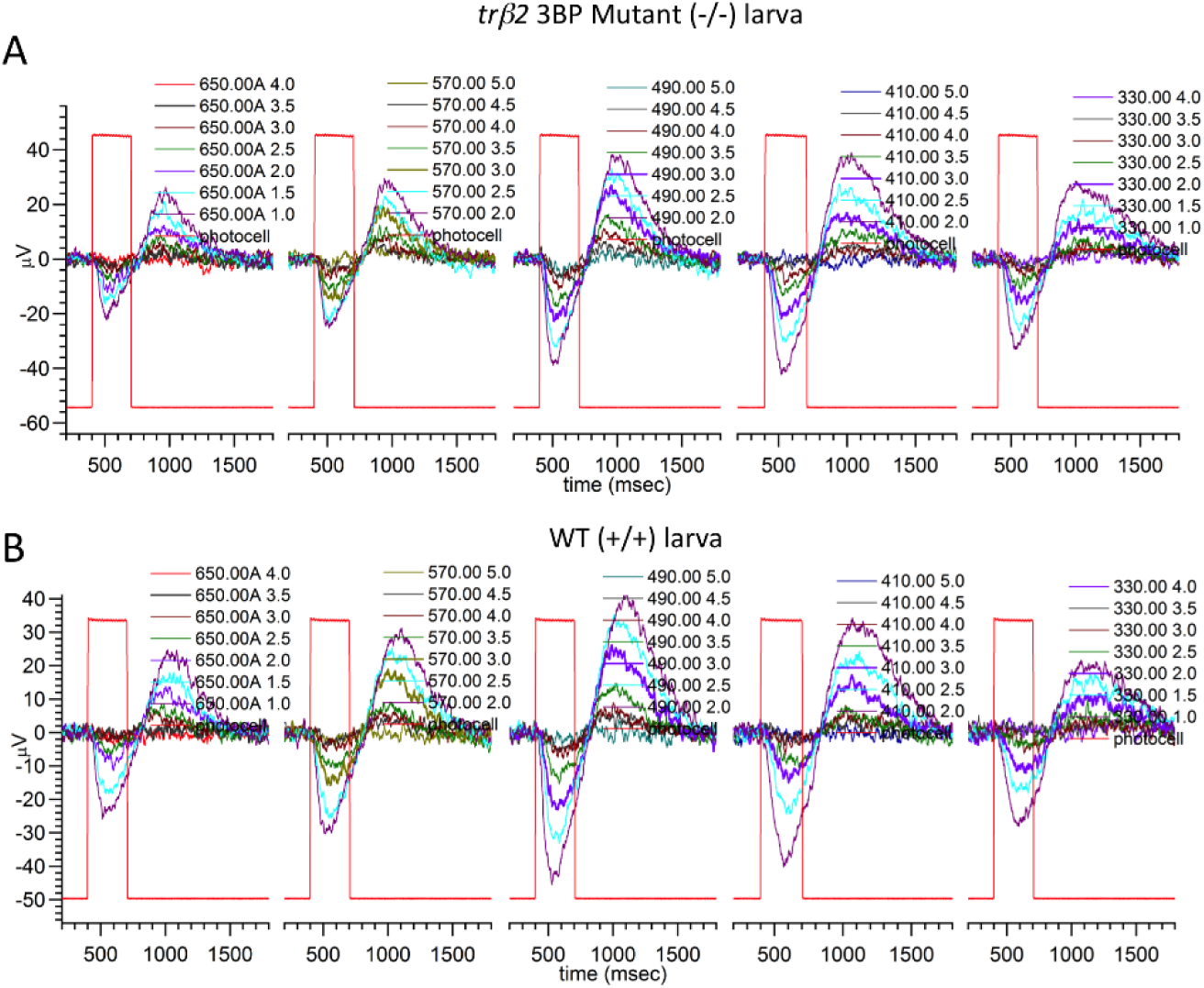
3BP larval cone PIII ERG spectral response traces. (A) The 3BP mutant larva is able to respond to all wavelengths from 650-330 nm, with no significant differences from the wild type control (B). Larvae are 5-7 dpf.

One potential action of a retinal mutation is to change the maximal voltage amplitudes of PIII ERGs. Table 1 gives the means and SEs of maximal dataset amplitudes in six genetic groupings for the *trβ2* gene. In the *6BP*+*1* and *3BP* rows are the results for 5-7 dpf larvae. For each mutation, larvae are from the same set of spawns so that other genetic differences are reduced. While there is a trend towards higher amplitude responses in the *6BP*+*1* mutants and heterozygous mutants, the trend did not achieve significance [ANOVA, F(2, 33) = 1.74, *P* = 0.19]. No amplitude changes were noted in the *3BP* strains [ANOVA, F(2, 61) = 1.80, *P* = 0.17]. Since amplitude changes are not significant, datasets from the larval genetic treatment groups were normalized to dataset maximal amplitudes and combined into treatment-level datasets for fitting to the spectral model. The number of amplitude-irradiance-wavelength points in these treatment level datasets varied from 560 for the *6BP*+*1* WT group to 2310 for the *3BP* heterozygous mutant group. As larvae were sequenced after physiological recording, while there was no prior knowledge of genotype, it was not possible to match the sizes of the treatment-level datasets in the 6 genetic groups.

**Table 1.** Maximal dataset amplitudes in *trβ2* genetic strains.

| *trβ2* strain | age | Units | WT (+/+) | Het mutant (+/-) | Mutant (-/-) | *P* |
| --- | --- | --- | --- | --- | --- | --- |
| *6BP+1* | 5-7 dpf | μV max | 29.4 ± 6.0 (8) | 57.4 ± 10.1 (16) | 76.4 ± 23.7 (12) | 0.19 |
| *3BP* | 5-7 dpf | μV max | 37.4 ± 3.9 (14) | 30.3 ± 2.0 (33) | 37.2 ± 4.8 (17) | 0.17 |
| *6BP+1* | 8-18 mo | μV max | 8.5 ± 1.2 (11) | 8.7 ± 1.2 (9) | 6.6 ± 0.7 (12) | 0.20 |
| *3BP* | 8-18 mo | μV max | 8.5 ± 1.2 (11) | 14.0 ± 1.9 (10) | lethal | * |

Amplitudes are means and SEs, with the number of datasets in parentheses. The *P* statistic is ANOVA probability

Model fits for irradiance response functions in 6 treatment level datasets are seen in Figure 9. Points are means from within the treatment level-datasets, while the curves result from a model fit to the entire dataset. These are not single-cone irradiance response curves, but the summation of multiple cone signals at each wavelength. The model curves fit all the six different larval genetic datasets well. The most severely altered irradiance response patterns arise in the *6BP*+*1* -/- mutant (Fig. 9C). Responses to red 650 nm wavelengths remain at near ‘0’ amplitude, regardless of brightness. The 490 nm curve is lowered in amplitude, and the 410 nm curve is displaced away from the 370 nm curve towards brighter irradiances as compared to WT (Fig. 9A). This shift appears in lesser degree in the *6BP*+*1* heterozygous mutant (Fig. 9B). The irradiance response patterns for *3BP* homozygous mutant (Fig. 9F) and heterozygous mutant (Fig. 9E) are qualitatively like those of WT larvae from the same spawns (Fig. 9D).

**Fig 9:**
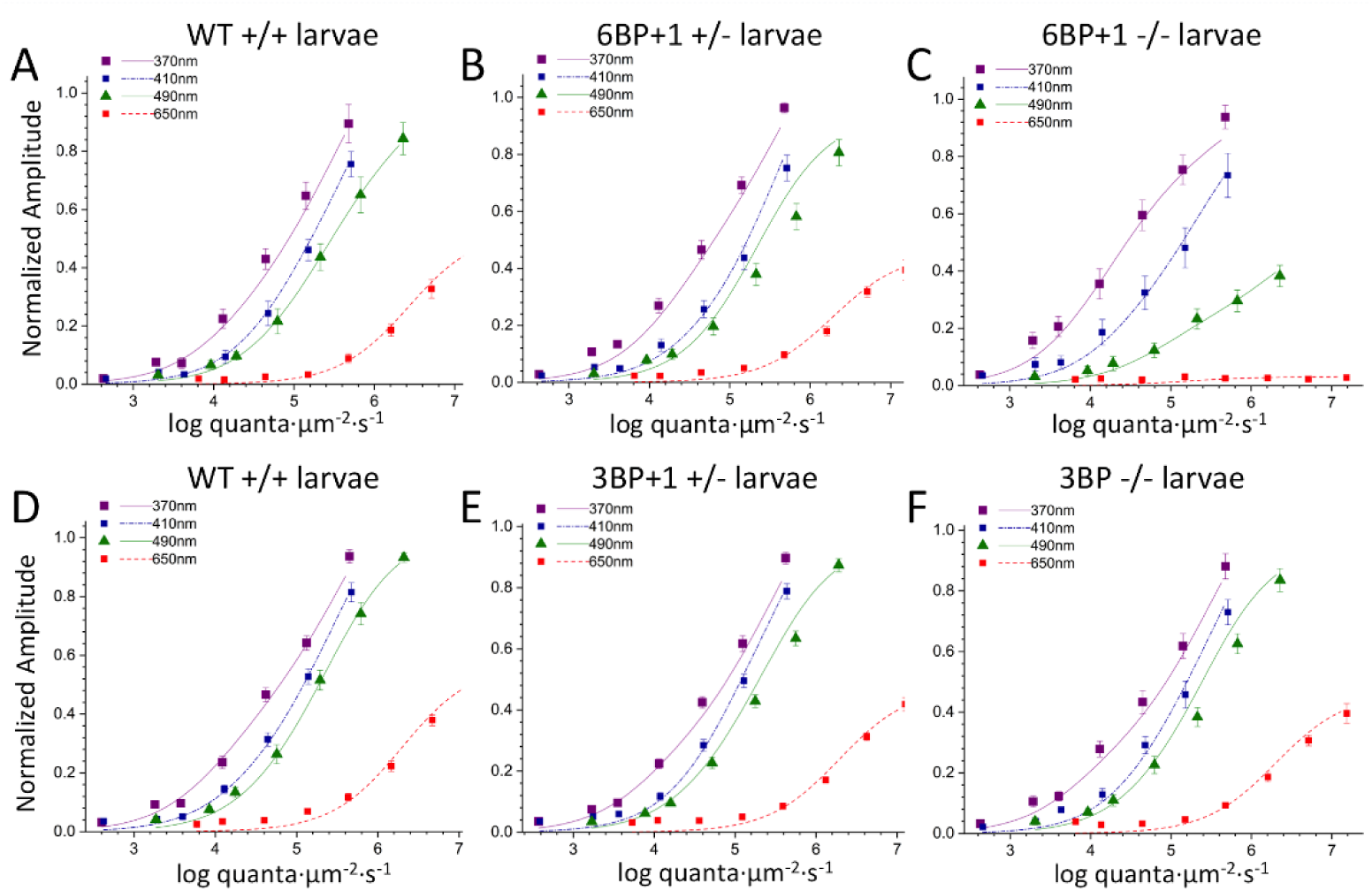
Irradiance plots: change in amplitude of cone PIII ERG responses as a function of brightness level. (A, B) The wild type and *6BP*+*1* heterozygote plots are similar, while the *6BP*+*1* mutant (C) shows a significant loss of response to 650 nm that does not change at higher irradiance levels. (D-F) Wild type, *3BP* heterozygote, and *3BP* mutant larvae do not significantly differ in their respective irradiance plots.

Spectral curves, which are the amplitudes at specific wavelengths normalized relative to the maximal amplitude for all wavelengths, are model generated in Figure 10. Response amplitudes are calculated for a constant quantal level of 4.6 log(quanta·µm^-2^·s^-1^), which was below semi-saturation for all but UV cone types. In the *6BP*+*1* mutant fish (Fig. 10A) amplitudes of response are greater than WT for wavelengths shorter than 420 nm, and less than WT for wavelengths longer than 420 nm. The *6BP*+*1* heterozygous mutant amplitude is slightly increased for wavelengths shorter than 410 nm. In the *3BP*, p.Tyr61del larvae, spectral curves are remarkably similar for mutants, heterozygotes and WT (Fig. 10B).

**Fig 10:**
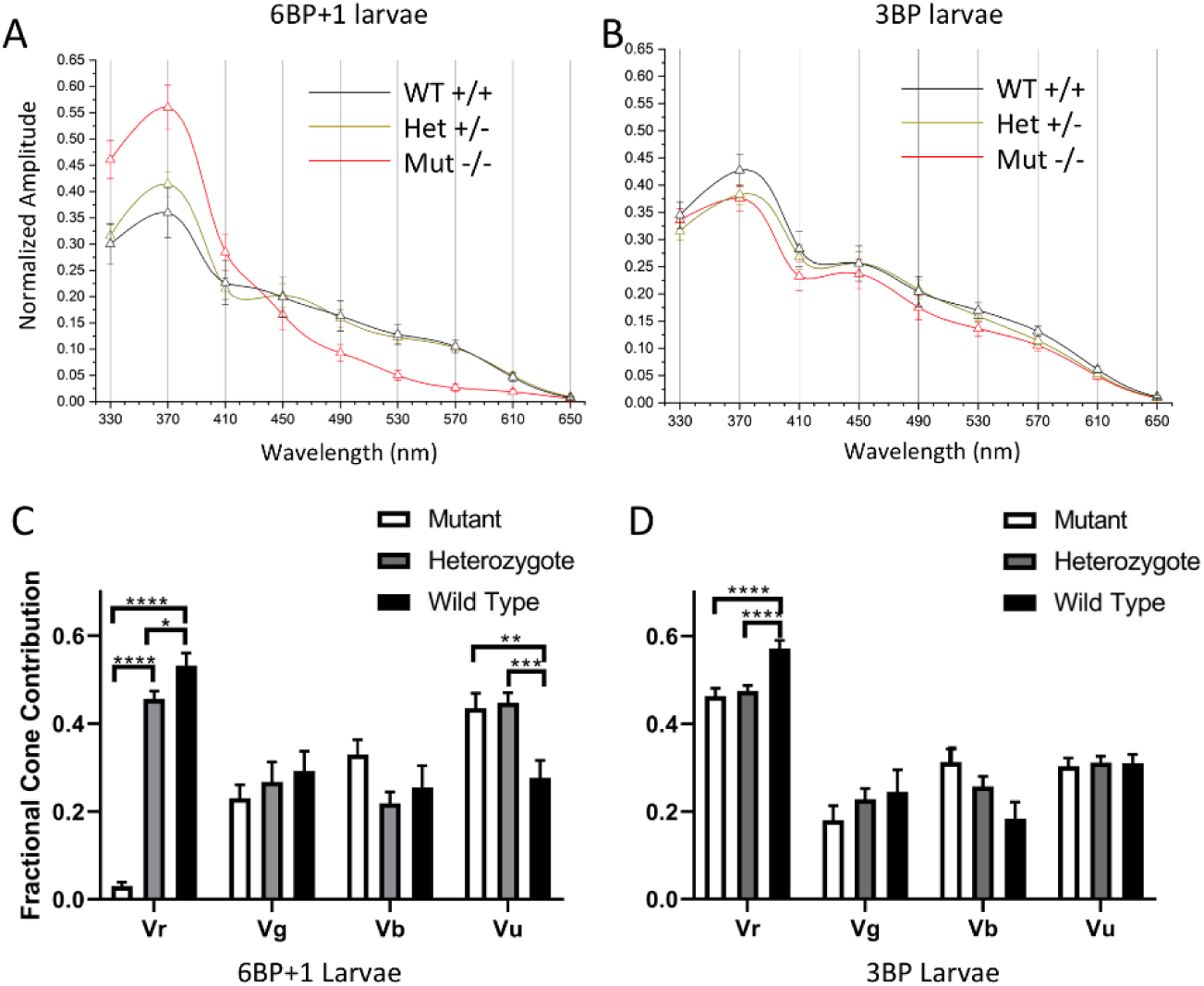
Spectral plots and cone contribution comparisons for *6BP*+*1* and *3BP* larva. (A) Shifting from 330 nm to 650 nm the *6BP*+*1* mutant shows a large drop in response amplitude for long wavelengths and a strong preference for UV wavelengths compared to the wild type and heterozygote. (B) The *3BP* spectral plot shows a similar spectral pattern between all genotypes in response amplitude across the wavelengths. (C) The *6BP*+*1* mutant displays a significant drop in red cone contribution (Vr) accompanied by an increase in UV cone contribution (Vu). (D) The cone contribution comparisons between the *3BP* genotypes show a significant decrease in Vr for the mutant and heterozygote.

### Zebrafish larvae with the 6BP+1 trβ2 mutation have a strong cone-signal phenotype

The goal of the model is to rationalize spectral datasets in terms of changes in the parameters of responsiveness for cones of different spectral types. *Vmax*, the cone saturation amplitude, is a model-fit index proportional to the number of cones and their response strength as individuals. *Vmax* values for red, green, blue, and UV cones (Vr, Vg, Vb, and Vu) are compared in Figure 10C and D. As expected from the above results, the red cone amplitudes (Vr) in the *6BP*+*1* -/- mutant were eliminated (Fig. 10C) [ANOVA, F(2, 2517) = 205.8, *P* = 0.0000]. The red cone fractional contribution in the mutant (0.030 ± 0.008) was significantly decreased in comparison to both WT (0.532 ± 0.028) and heterozygous mutant (0.456 ± 0.018) (Fig. 10A, Tukey post hocs, *P* = 0.0000). In addition, there was a significant red-cone amplitude-reduction phenotype for the heterozygous mutant in respect to WT (Tukey post hoc, *P* < 0.05). Among the *6BP*+*1* larval siblings, the three genetic outcomes of the spawns showed UV cone (Vu) amplitudes that were significantly impacted [ANOVA, F(2, 2157) = 7.9, *P* < 0.001]. There were increased amplitudes of UV cone signals in both mutant and heterozygous mutant larvae as compared to WT (Tukey post hocs, P < 0.01, P < 0.001, respectively, Fig 10A). The fractional amplitudes were 0.276 ± 0.040 (WT), 0.447 ± 0.022 (heterozygous mutant), and 0.435 ± 0.034 (mutant). No significant differences among the larval sibling genetic types were found for green or blue cone amplitudes [ANOVA: Vg, F(2, 2517) = 0.38, *P* = 0.38; Vb, F(2, 2517) = 0.48, *P* = 0.61]. Results suggest a significant role for the *trβ2* gene in the apportionment of signal strength among red, and UV cones but not green or blue cones.

### Zebrafish larvae with the 3BP trβ2 mutation have a weak cone-signal phenotype

The irradiance-response (Fig. 9D-F) and spectral plots (Fig. 10B) for the *3BP* deletion mutant do not suggest a spectral phenotype, however the modeled larval cone component analysis indicates a subtle change. For the *3BP* deletion mutant, heterozygous mutant, and WT larvae, red cone amplitudes (Vr) varied significantly [ANOVA, F(2, 4476) = 10.76, *P* = 0.0000].The red cone fractional contributions in the *3BP* mutant (0.463 ± 0.018) and the heterozygous mutant (0.474 ± 0.013) were significantly less than WT (0.573 ± 0.017) (Fig. 10D, Tukey post hocs, *P* < 0.0001). No other cone type varied significantly among the *3BP* WT, heterozygous mutant, and mutant larvae [ANOVA: Vg, F(2, 4476) = 0.88, *P* = 0.415; Vb, F(2, 4476) = 2.59, *P* = 0.075; Vu, F(2, 4476) = 0.327, *P* = 0.721]. The trend towards increased blue-cone signal (Vb, Fig. 10B) in the *3BP* heterozygote as compared to WT, while not significant (Tukey, *P* = 0.062), is also found in the *6BP*+*1* mutation. The *3BP* protein appears at least partially competent in directing progenitors into the red-cone pool.

### Cone sensitivity peaks in larval trβ2 genetic strains

The trβ2 nuclear receptor is a transcription factor, and a candidate molecule to regulate cone opsin expression. For this reason, we compare the fit values for cone sensitivity peaks in the *6BB*+*1* and *3BP* mutants or heterozygous mutants to WT siblings (Table 2). In no case where a red cone sensitivity peak could be detected was it significantly changed by either mutant or heterozygous mutant genetics. All peaks lay in a tight range between 554 and 557 nm, similar to previous reports for WT larvae, and consistent with *LWS2* expression [16]. The missing peak, of course was the *6BP*+*1* mutant, where there is no red-cone signal. In the *6BP*+*1* mutant and heterozygous mutants green, blue and UV cone peaks were unaffected (Table 2) and in a range reported for WT larvae [16], and in a range corresponding to G1 (*RH2-1*) for green cones, B1 (*SWS2*) for blue cones, and U (*SWS1*) for UV cones. Because of low signal amplitude in larvae, the inability to find green cone sensitivity peaks in *6BP*+*1* mutants or the entire cohort of *3BP* variants is not taken as an indicator of gene action.

**Table 2.** Peak sensitivities of larval cones in *trβ2* genetic strains.

| Larval *trβ2* strain | Cone type | Unit s | WT (+/+) | Het mutant (+/-) | Mutant (-/-) | *P* |
| --- | --- | --- | --- | --- | --- | --- |
| 6 BP+1 | red (LWS) | nm | 553.9 ± 3.1 (560) | 556.7 ± 2.8 (1120) | no fit | 0.54 |
|  | green (Rh2) | nm | 460.2 ± 7.1 (560) | 466.4 ± 4.3 (1120) | no fit | 0.43 |
|  | blue (SWS2) | nm | 401.7 ± 8.7 (560) | 396.4 ± 5.2 (1120) | 399.7 ± 4.4 (840) | 0.81 |
|  | UV (SWS1) | nm | 352.7 ± 4.2 (560) | 350.3 ± 3.1 (1120) | 354.6 ± 2.7 (840) | 0.61 |
| 3 BP | red (LWS) | nm | 555.5 ± 2.0 (980) | 556.0 ± 1.9 (2310) | 555.2 ± 2.8 (1189) | 0.97 |
|  | green (Rh2) | nm | no fit | no fit | no fit | ----- |
|  | blue (SWS2) | nm | 420.2 ± 6.3 (980) | no fit | 399.0 ± 3.8 (1189) | ** |
|  | UV (SWS1) | nm | 361.3 ± 2.0 (980) | 357.5 ± 2.0 (2310) | 347.4 ± 3.6 (1189) | ** |

Larval cone wavelength peaks are fit to treatment-level datasets using equation 3. Values are wavelength peaks ± SEs (n), where n is the number of amplitude-irradiance-wavelength points fit in each treatment-level dataset.

In the *3BP* mutant, both blue (*SWS2*) and UV (*SWS1*) cone peaks shifted significantly to shorter wavelengths in the mutant genotype. A difference in blue-cone peaks has elsewhere been noted among different WT cohorts, and attributed to the presence of two opsins (B1 and B2) in the blue waveband [16]. These authors also noted that *roy orbison* mitochondrial membrane protein mutation [17] altered the UV cone spectral peak.

### 6BP+1 mutants do not show an optomotor response to a red and black striped stimulus

Zebrafish larvae tend to swim in the direction of a drifting grating. This is called the optomotor response (OMR). The response is reported to be driven by both red and green sensing cones in larvae [18], but dependent on red cones in adults [19]. We tested whether this behavior was lost in *6BP*+*1 trβ2* mutants. Larvae of different genotypes were placed in a tissue-culture plate atop of laptop monitor, where the black/white grating was presented for one minute. The plate was divided into 3 regions (Fig. 11A) and the number of larvae in regions 1 and 3 compared before and after stimulation using a χ^2^ test. For WT larvae (Fig. 11A, 11A’), there were 8 larvae in region 1, the top half of the plate, before the stimulus, but only 3 afterwards. In region 3, the bottom rim of the plate, there were 0 larvae before the stimulus, but 10 larvae after the stimulus. The change in distribution is significant (χ^2^, *P* < 0.00001). Both *6BP*+*1* heterozygous (Fig. 11B, 11B’) and homozygous mutant (Fig. 11C, 11C’) genotypes showed significant OMR behavior (χ^2^: +/-, *P* < 0.001; -/- *P* < 0.00001) to the black/white grating. The *6BP*+*1* mutants are capable of vigorous of OMR visual behavior even in the absence of physiological signals from red cones.

**Fig 11:**
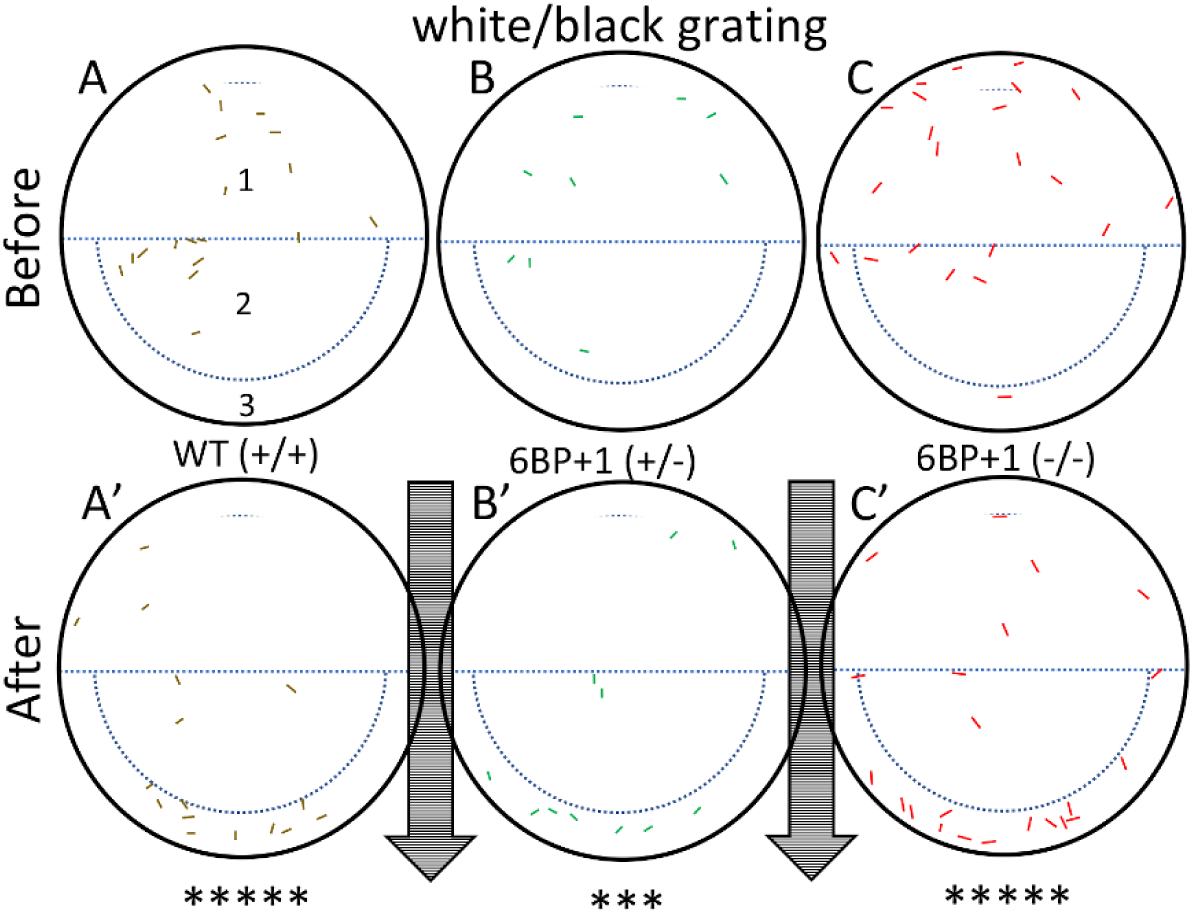
Optomotor response test. Controls use a black and white grating pattern with wild types (+/+), *6BP*+*1* heterozygotes (+/-), and *6BP*+*1* mutants (-/-) at 7 dpf. Colored markings in the dishes represent larval zebrafish locations before and after the one minute stimulus. The arrows indicate the direction of the stimulus. (A, B, C) Wild types, heterozygotes, and mutants respectively start scattered in the dish. (A’, B’, C’) Each genotype moved with the stimulus showing significant pooling of larvae at the bottom edge.

Because of a lack of LWS cone physiology, OMR responses to red/black drifting gratings were not expected in *6BP*+*1* homozygous mutants, and in fact were not found (Fig 12). Of 16 larvae in region 1 of the mutant plate (Fig. 12C) 16 remained after stimulation with a red/black grating (Fig. 12C’). In region 3, there was 1 larva before and after stimulation. These results are consistent with a lack of mutant OMR for red/black gratings (χ^2^, *P* = 1). WT and heterozygous mutant larvae (Figs. 12A, 12A’; 12B, 12B’) showed significant OMR behavior under the same conditions (χ^2^: +/+, *P* < 0.05; +/- *P* < 0.05). These genotypes show that red cone signals in themselves are sufficient for OMR in wildtype fish but the absence of red cones prevents a response in the *6BP*+*1* mutant fish.

**Fig 12:**
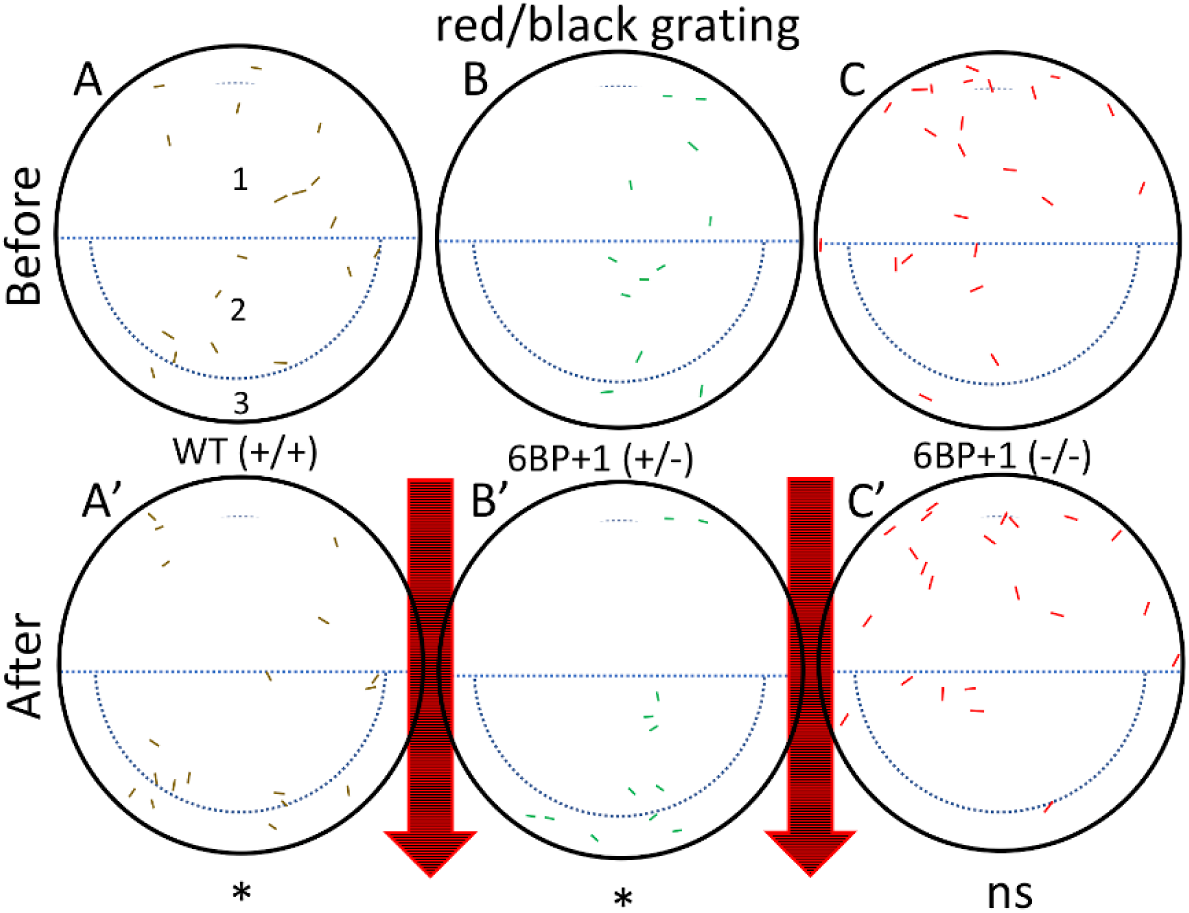
Optomotor response test using a red and black stimulus. The wild type (A, A’) and heterozygote (B, B’) larva moved with the stimulus as seen in the control test. (C, C’) The mutant larvae did not move with the stimulus and remained scattered.

There was no significant difference between the wild types, *3BP* het mutant, and *3BP* mutants for OMR. Each genotype had an OMR to every stimulus (data not shown).

### 6BP+1 mutant fish have a significantly lower optokinetic response with red/black stimulus (610 nm) and green/black stimuli (550 nm)

Larval zebrafish eyes track rotating patterns, and then snap back to reset [20]. This optokinetic response (OKR) is an obligatory visual reflex among vertebrates. The question is the role of red cones in this reflex. In Figure 13 eye movements are counted for colored rotating windmills presented on a laptop monitor for one minute. Mutant *6BP*+*1 trβ2* homozygous (-/-) alleles, WT (+/+), and heterozygous mutants (+/-) were shown white/black, red/blue, red/black, and green/black pinwheels (Fig. 13A, A’; B, B’; C, C’; D, D’). The *6BP*+*1* homozygous mutant fish (-/-) show OKR responses only for white/black and red/blue pinwheels, but not red/black or green/black pinwheels. The mutant red/black OKR responses (Mdn = 0.0) were significantly lower than WT (Mdn = 1.0) (*Mann Whitney,* U = 188, *P* < 0.05) and the heterozygous mutant (Mdn = 2.0) (*Mann Whitney,* U = 197, *P* < 0.01, Fig. 13C’). There was a significant decrease in the number of green/black OKRs between the homozygous mutant fish (Mdn = 0.0) and WT (Mdn = 2.0) (*Mann Whitney*, U = 168, *P* < 0.01) or heterozygous mutant (Mdn = 3.0) (*Mann Whitney*, U = 118, *P* < 0.0001. Fig. 10D’). In respect to OKR behavior, the *6BP*+*1 trβ2* mutant fish do not respond to the monitor colors red or green.

**Fig 13:**
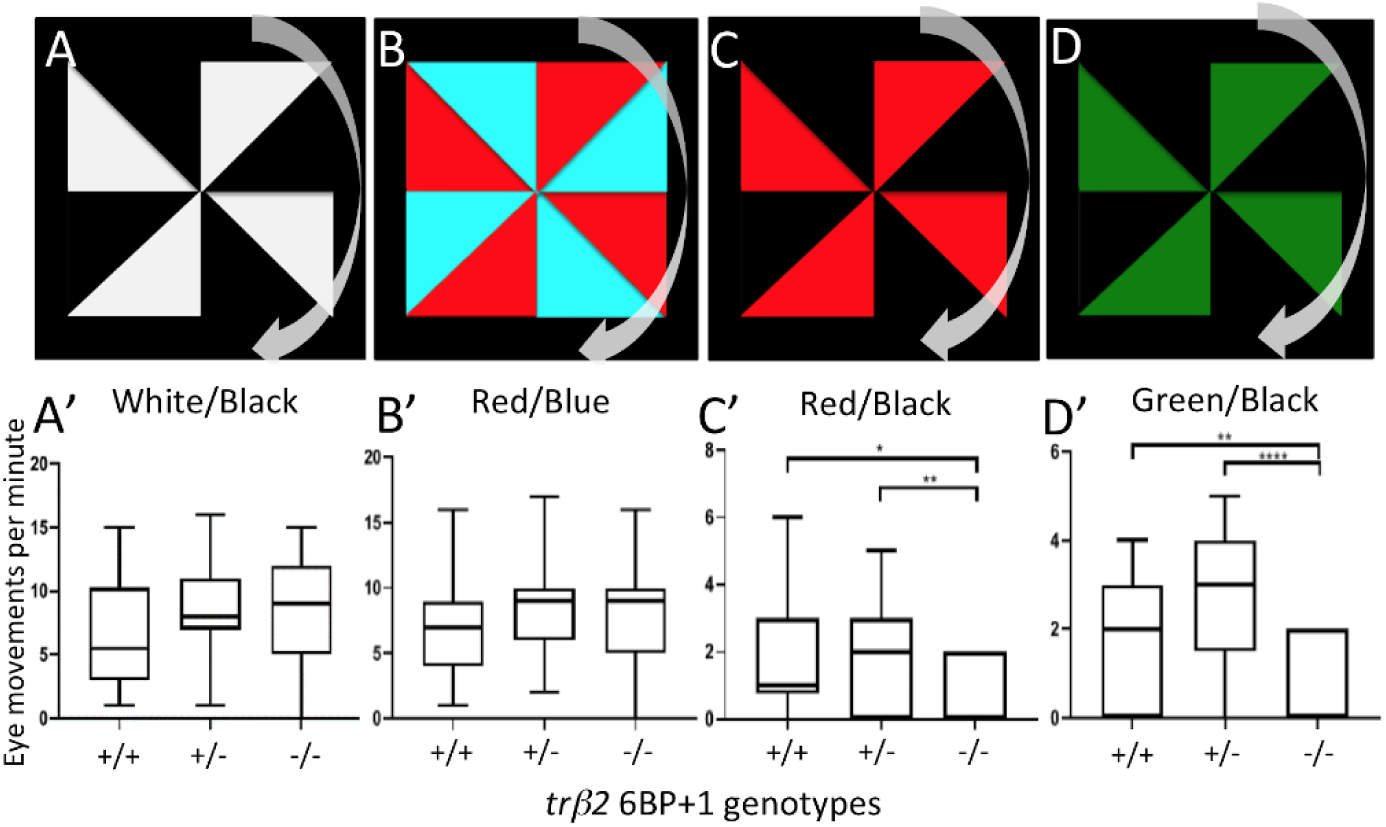
Optokinetic response tests. (A, B, C, D) Each larva at 9 dpf was exposed to these four stimuli in this respective order. The pinwheel patterns moved clockwise. (A’, B’) The wild type, 6BP+1 heterozygote, and 6BP+1 mutant responded similarly to the white/black and red/blue stimuli. (C’, D’) The mutant had significantly less eye movements in response to the red/black and green/black stimuli.

### The loss of red response in the 6BP+1 mutant larvae is maintained in adult zebrafish

As in *6BP*+*1* homozygous mutant larvae, adult *6BP*+*1* mutant fish are unresponsive to long wavelength stimuli. In a cone-PIII dataset from an adult *6BP*+*1* mutant fish (Fig. 14A), no cone PIII responses are evoked by 650 nm stimuli of any brightness, whereas WT and heterozygous mutants respond well at multiple brightness’s (Fig. 14B, 14C). For the *6BP*+*1* mutant only the brightest of 570 nm stimuli produces a response, while the WT and *6BP*+*1* heterozygous adults respond vigorously to multiple stimulus irradiances.

**Fig 14:**
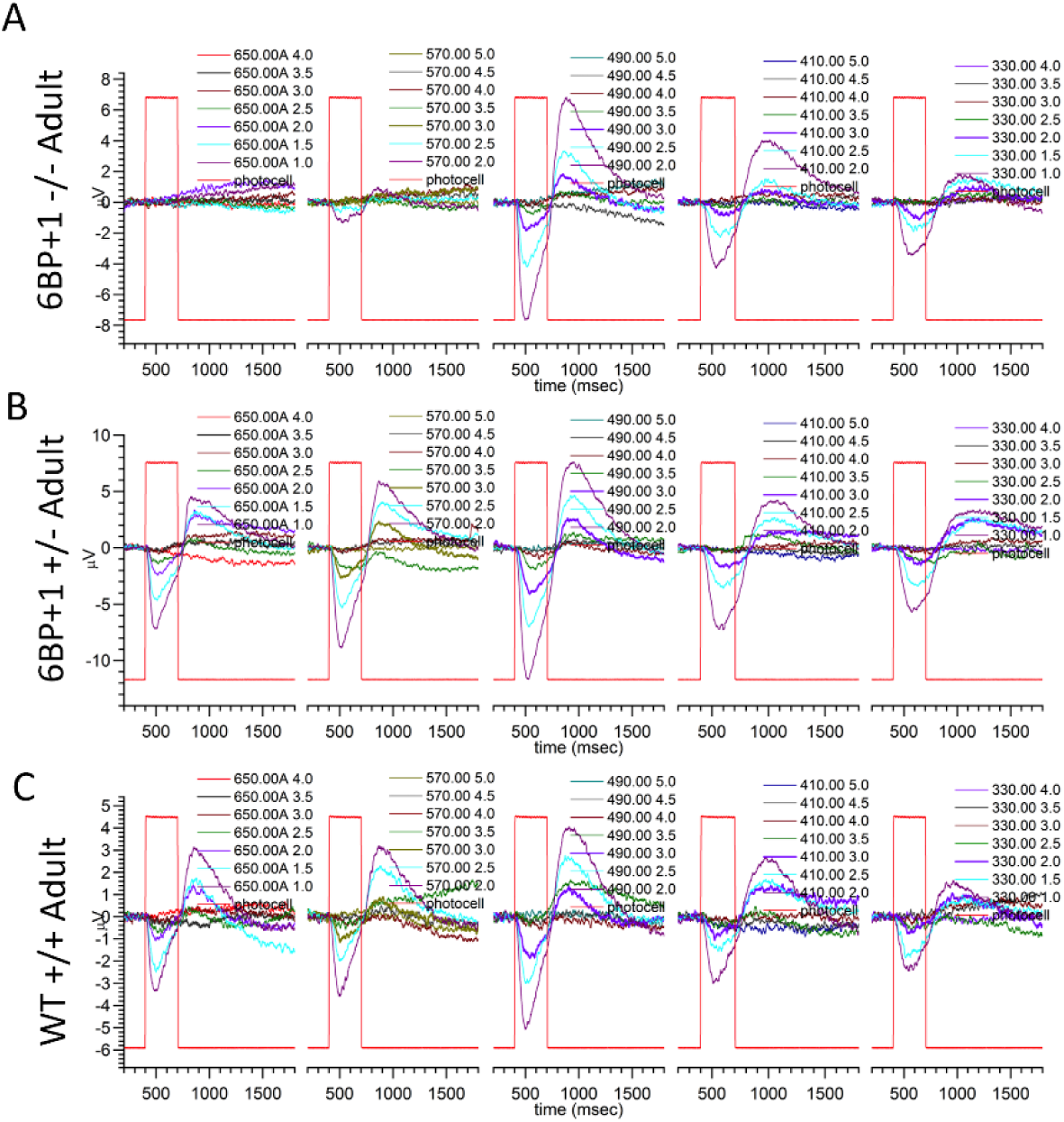
Adult PIII ERG response plots. (A) 6BP+1 mutant adults did not respond to 650 nm, but did respond to the maximum brightness level of 570 nm. (B) Heterozygotes respond to every wavelength. (C) Wild types respond to all wavelengths.

For adult genotypes, the treatment-level irradiance-response, spectral, and cone component characteristics are summarized in Figure 15. For WT the model is fit to 770 amplitude-irradiance-wavelength points (7 eyecups), for the *6BP*+*1* heterozygous mutant, 630 points (6 eyecups) and the *6BP*+*1* mutant, 840 points (7 eyecups). Dataset maximal amplitudes were not affected by genotype (Table 3).

**Table 3.**
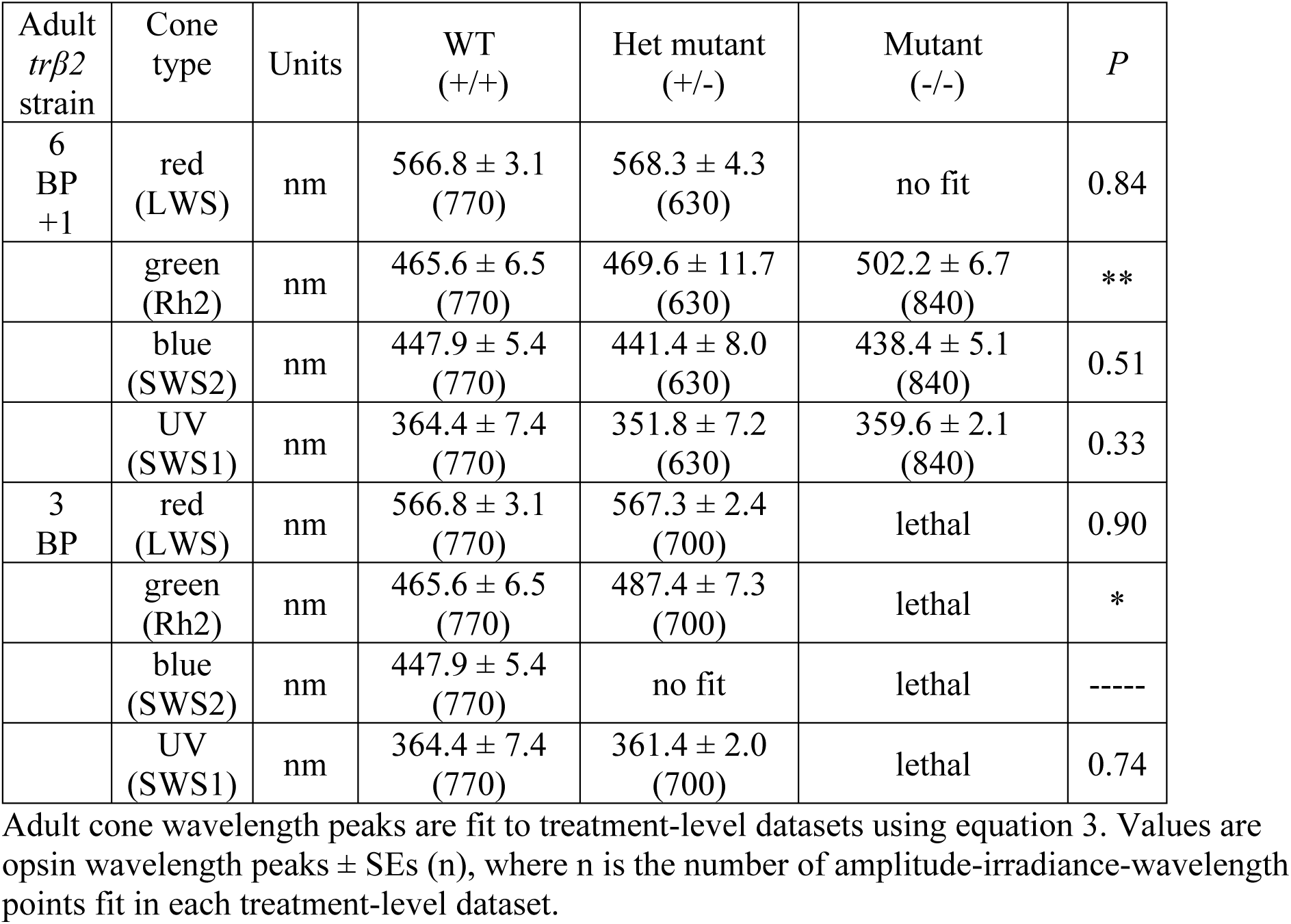
Peak sensitivities of adult cones in *trβ2* genetic strains.

| Adult *trβ2* strain | Cone type | Units | WT (+/+) | Het mutant (+/-) | Mutant (-/-) | *P* |
| --- | --- | --- | --- | --- | --- | --- |
| 6 BP +1 | red (LWS) | nm | 566.8 ± 3.1 (770) | 568.3 ± 4.3 (630) | no fit | 0.84 |
|  | green (Rh2) | nm | 465.6 ± 6.5 (770) | 469.6 ± 11.7 (630) | 502.2 ± 6.7 (840) | ** |
|  | blue (SWS2) | nm | 447.9 ± 5.4 (770) | 441.4 ± 8.0 (630) | 438.4 ± 5.1 (840) | 0.51 |
|  | UV (SWS1) | nm | 364.4 ± 7.4 (770) | 351.8 ± 7.2 (630) | 359.6 ± 2.1 (840) | 0.33 |
| 3 BP | red (LWS) | nm | 566.8 ± 3.1 (770) | 567.3 ± 2.4 (700) | lethal | 0.90 |
|  | green (Rh2) | nm | 465.6 ± 6.5 (770) | 487.4 ± 7.3 (700) | lethal | * |
|  | blue (SWS2) | nm | 447.9 ± 5.4 (770) | no fit | lethal | ----- |
|  | UV (SWS1) | nm | 364.4 ± 7.4 (770) | 361.4 ± 2.0 (700) | lethal | 0.74 |

Adult cone wavelength peaks are fit to treatment-level datasets using equation 3. Values are opsin wavelength peaks ± SEs (n), where n is the number of amplitude-irradiance-wavelength points fit in each treatment-level dataset.

**Fig 15:**
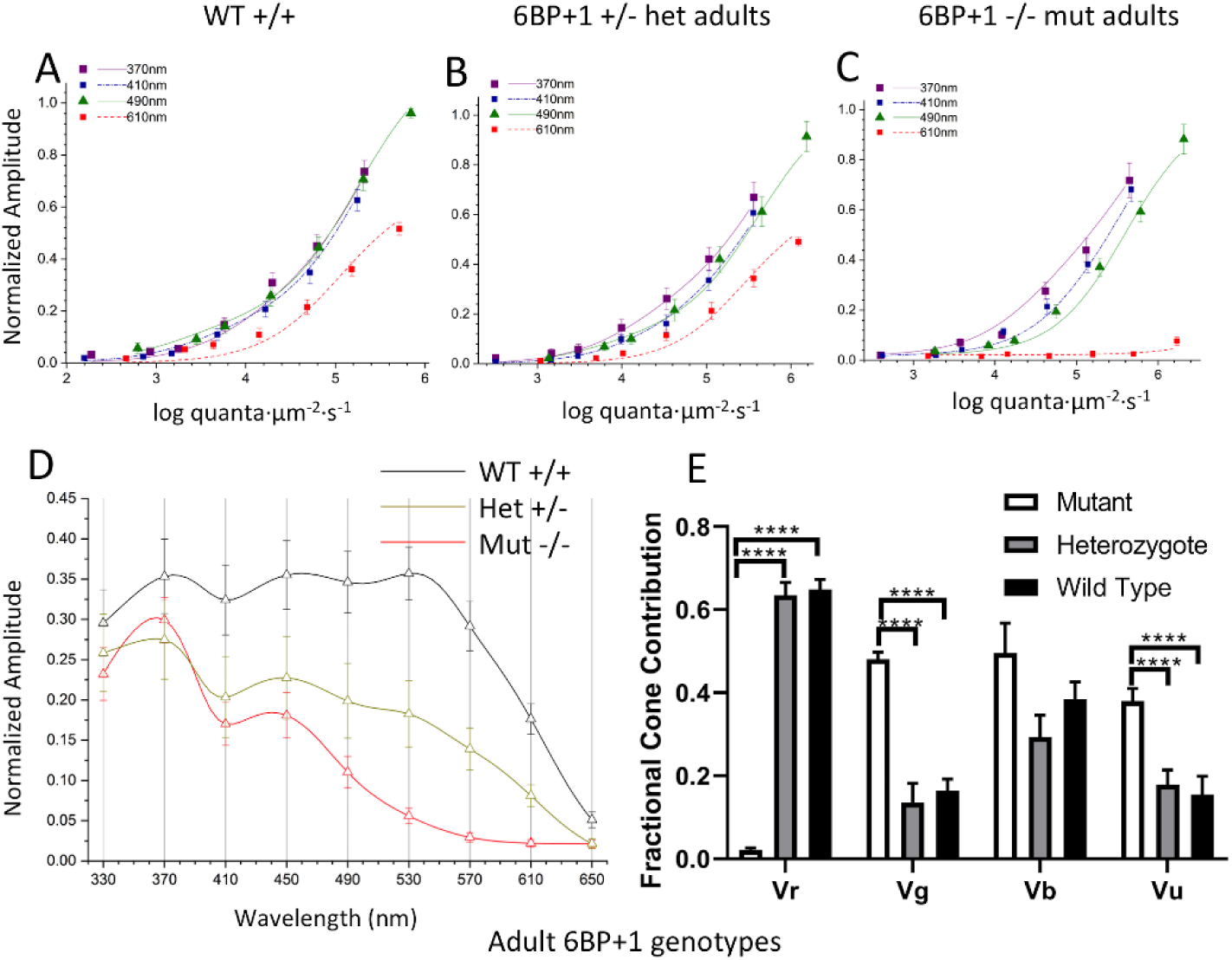
Spectral analysis of 6BP+1 adults. (A, B, C) The irradiance plots for each genotype showed the loss of response to 610 nm in the mutant regardless of brightness. (D) The spectral plot showed a significant loss of amplitude in the heterozygote and mutant at long wavelengths compared to the WT. (E) The mutant shows loss of Vr, increase in Vg, and increase in Vu.

The model curves closely fit irradiance response data points at multiple wavelengths in the three genotypes (Fig. 15A, B, C). In the *6BP*+*1* mutant (Fig. 15C), there is a complete loss of responsiveness at 610 nm, whereas WT and heterozygous mutants show increasing response amplitudes with brighter 610-nm irradiances (Fig. 15A, 15B). The 370 nm, 410 nm and 490 nm curves are closely bunched for WT, but separate for *6BP*+*1* heterozygotes and especially mutants, indicative of spectral differences. The spectral curves, PIII fractional amplitudes for constant quantal irradiance across the spectrum, show a different spectral phenotype for each genotype (Fig. 15D). The stimulus is 4.6 log(quanta·µm^-2^·s^-1^), in the range of semi-saturation for cone types. While amplitudes for UV stimulation are similar in WT, *6BP*+*1* mutant and heterozygote, the mutant and heterozygote evoke lesser amplitudes at longer wavelengths. In the 6PB+1 mutant, there is no spectral response for wavelengths longer than 570 nm, and the heterozygote response at these wavelengths is intermediate compared to WT.

In the adult *6BP*+*1* mutant genotype, red and UV cone saturation amplitudes (Vr, Vu, Fig. 15E) are altered in a way similar to *6BP*+*1* larvae (Fig. 10C). Vr is severely diminished in the adult *6BP* +*1* mutant fish as compared to either WT or heterozygous mutants [ANOVA: F(2, 2237) = 312.8, *P* = 0.0000]. Unlike *6BP*+*1* mutant larvae, heterozygous adults show no significant diminution in Vr as compared to WT. In the adult *6BP*+*1* mutant fish Vu is significantly greater than WT [ANOVA: F(2, 2237) = 11.7, *P* = 0.0000]. Unlike *6BP*+*1* larvae, the adult Vu value for heterozygotes is not increased, but the same as WT. The greatest departure from the larval pattern is in the green cones. The adult *6BP*+*1* mutant Vg amplitude is greatly increased, both in respect to adult WT and to adult heterozygotes [Fig. 15E: ANOVA: F(2, 2237) = 13.0, *P* = 0.0000]. The missing red cone contribution is distributed across green, blue, and UV cones in the mutant.

### Cone sensitivity peaks in adult trβ2 genetic strains

We compare the fit values for cone sensitivity peaks in the *6BP*+*1* and *3BP* mutants or heterozygous mutants to WT siblings (Table 3). In no case where red cone sensitivity peak could be detected was it significantly changed by the presence of the mutant gene. All peaks lay in a tight range between 566 and 569 nm, similar to previous reports for WT adults, and consistent with adult *LWS1* expression [16]. There are missing peaks in the *6BP*+*1* and *3BP* mutant, in the first mutant because of a lack of red-cone signal, and in the second because the mutation is embryonic lethal. In the *6BP*+*1* mutants and heterozygous mutants blue and UV cone peaks were unaffected (Table 3) and in the *3BP* heterozygote, the UV cone peak was unaffected. In *6BP*+*1* mutant the green cone peak was significantly shifted to longer wavelengths. At 502 nm this might represent expression of *RH2-4* [11]. Both the wavelength and sensitivity change of green cones in the *6BP*+*1* mutant suggest an alteration of the mutant’s green cones. A significant long-wavelength shift of the adult *3BP* heterozygote green cone is also noted. The semi-saturation kg value in the *6BP*+*1* mutant fish (4.76 ± 0.11) was increased compared to WT (3.17 ± 0.17). The change was significant (*t* = 7.58, df = 1468, *P* < 0.0001). The suspiciously similar actions of *6BP*+*1* and *3BP* mutant *trβ2* alleles on green cone opsin peaks suggests a long term role for *trβ2* in green cone maturation.

## Discussion

Two *trβ2* mutant strains provide insights into the roles of this transcription factor in the vertebrate retina. Both mutations target the same PY amino acid sequence in the N-terminal region of the gene. This is a ‘transactivation’ region, critical to binding other transcription factors and to the activation or inhibition of the transcription process [5]. Both mutants significantly reduced function, but the phenotypes differed.

### Frameshift mutant

The loss of response to red wavelengths in the zebrafish *6BP*+*1 trβ2* mutant is consistent with the losses seen in mouse and human mutants, corroborating the conservation of *trβ2’s* function among vertebrates in mediating the development of long-wavelength cones and long-wavelength sensory pathways [1, 3, 4]. In addition to the significant decrease in responses to red wavelengths, the absence of *trβ2* in zebrafish retina development leads to an increase in UV cone response amplitudes and fractional UV cone contribution. This suggests a negative regulation of UV cones specifically by *trβ2*, since thyroid receptors are capable of both positive and negative regulation [5, 21]. In larval zebrafish, the significant decrease in, if not elimination of, Vr and increase in Vu for the *6BP*+*1* heterozygote in comparison to the wild type identifies a heterozygous phenotype resembling the mutant but to a lesser degree.

Elements of the larval cone phenotype are retained in adult zebrafish, but in addition there are actions on green cones. Though we do not see any significant changes in green cones in larvae, we see a significant increase of Vg in the *6BP*+*1* adult mutant, a decrease in green-cone sensitivity (kg), and a shift to a longer-wavelength spectral peak for green cones. Of the four *RH2* (green opsin) subtypes, *RH2-2* has the highest expression level in wild type adult zebrafish retina with an in-solution spectral peak of 476 nm [11, 22], and *in situ* spectral measurements corroborate a peak at ∼480 nm [16]. In our *6BP*+*1* mutant adult we see the green opsin spectral peak is fit to 502 nm, which is closely aligned with the *RH2-4* solution peak at 505 nm suggesting the loss of trβ2 consequently induced an increase of *RH2-4* expression [11]. The long-wavelength shift in mutant green-cone opsin is a partial spectral compensation for red-cone loss.

In development there is a shift from LWS2 to LWS1 opsin [23] and a concomitant shift in physiological red-cone opsin peak from 556 nm to 572 nm [16]. Thyroxin T3 mediates this shift [24]. While *6BP*+*1* mutant fish lack LWS opsins altogether, heterozygous mutant fish show a developmental shift in red-cone spectral peak similar to wild type, so that at least modest reductions in trβ2, while otherwise evidencing a spectral phenotype, do not impede the LWS opsin shift with development.

In the *6BP*+*1* mutant fish there is no immunoreactivity for LWS opsins, in either larvae or adults. It also appears there is no immunoreactivity for arrestin (arr3), as determined with the zpr-1 antibody, raised against mouse arrestin3 and often used to label zebrafish red-green double cones. Zebrafish double cones express arr3a [30]. This suggests *trβ2* is a requirement for the expression of LWS1, LWS2, and arr3a proteins. The transgenic reporter *trβ2*:tdTomato used to make red cones [10] is inactive in the *6BP*+*1* mutant fish. The *trβ2* promoter used in this transgene would appear also to require the *trβ2* transcription factor for activation. As might be expected of a factor critical to the generation of a neural type, trβ2 activates multiple molecular pathways. Opsin-promoter-based transgenic reporters for green, blue, and UV cones continue to be expressed in the *6BP*+*1* mutant fish, and reveal the distributions of these remaining cone types. While the density of these types increases in the mutant, the cone ratios are preserved. Mechanisms for generation of cone density ratios may not require functional red cones, or trβ2.

Arrestin quenches the activated state of opsins. Morpholino suppression of arr3a in zebrafish larvae delays the recovery of b-wave responses [30]. In *6BP*+*1* mutant adult fish, altered green cone signals are observed. One of the changes is decreased sensitivity, as measured by half-saturation irradiance. This might be a consequence of arrestin loss. It is not clear whether the trβ2 influence of arr3 or Rh2 opsin expression in green cones is cell autonomous, or non-cell-autonomous. One model of cone mosaic formation involves cell-cell interactions among cone progenitors [31].

### Tyrosine deletion mutant

While the *3BP* mutant strain did not lose its ability to respond to red wavelengths entirely, the single codon deletion led to a significant decrease in red cone contribution. The Tyr61 deletion located in the N-terminus lies within a potential hormone-independent transactivation domain [5]. Previously it has been noted that patients with resistance to thyroid hormone due to mutations in the *thrb* ligand binding domain lack eye dysfunction [5]. In zebrafish, larvae with ablated thyroid glands maintain the same levels of *LWS2* expression as the controls with a working thyroid gland [24]. These findings suggest that *trβ2* is capable of inducing gene expression independent of thyroid hormone [5]. Our results are in line with that hypothesis. Although mutants still respond to red light, the decrease in cone contribution suggests that this N-terminus located tyrosine is important for the full functionality of *trβ2*. Contrary to our *6BP*+*1* mutant, the *3BP* mutant did not survive to adulthood, so there does seem to be a consequence of this mutation that is essential for life [25]. The latest detected developmental stage is 12 dpf as recordings were conducted at that time. Beyond 12 dpf the mutation appears to be lethal as 0 of the 56 genotyped adult (>3 months post fertilization) offspring from a *3BP* heterozygous breed were mutants. This result suggests further investigation as to why this specific deletion could prove to be lethal, given that the *6BP*+*1* mutant fish survive to adulthood, and at what developmental stage does it become lethal.

### Behavior

To supplement the ERG results for our complete *trβ2* knockouts, the optokinetic and optomotor response behavior tests gave insight into brain function. Wild type fish will swim with the stimulus and congregate at one side of the petri dish [26, 19]. However, the *6BP*+*1* mutant fish did not follow in suit for the red/black stimulus. The lack of response to the red and black stimulus by the *6BP*+*1* mutant fish showed that this loss starting in the cones is extended all the way to the brain. The larvae are unable to perceive red light, thus eliminating their ability to discern red from black. For the optokinetic response, the green light emitted from the monitor peaks at 550 nm [27]. This is closer to the larval red opsin spectral peak of 556 nm than green opsin at 461 nm, so although we perceive the stimulus as green it actually activates red opsin in larval zebrafish [16]. As a result, we see an even larger significant difference between the genotypes with green stimuli because it excites the wild types and heterozygotes more than the red stimulus, while maintaining a loss of response from the mutants. The control white/black stimulus, as well as the red/blue in the optokinetic experiments, resulted in no differences between the three genotypes, so red cones are not vital to the larval OKR pathway provided that other cones are stimulated.

### Summary

Together with clear changes in spectral physiology, the antibody stains and the image analysis of the adult *trβ2* chimera fish proved that *trβ2* is required for the differentiation of red cones and LWS opsin in zebrafish [1, 10, 28]. The larval stains of the *6BP*+*1* mutant showed a complete loss of red opsin, therefore a loss of functional trβ2 results in a loss of red opsin. The adult retina stains confirmed that the loss of red opsin in the mutant is maintained through development. The adult chimeric further confirmed the loss of red cones accompanied by additional structural changes. In adult zebrafish, the cones are arranged in alternating rows of ultraviolet/blue and red/green, which create a mosaic pattern [29, 15]. The loss of red cones in the mutant led to significant increases of the other three cone types, but they maintain their respective mosaic ratios as seen in the wildtype. These images align with the adult cone spectral-physiology as there was a significant increase in the green, blue, and UV cone PIII ERG contributions in *6BP*+*1* mutant fish. In mosaic patterning, unexpectedly, blue cones were in close proximity to green cones when red cones were missing, but the cones still attempt to form alternating blue/UV and green cone rows. This suggests the loss of *trβ2* and subsequently red cones leads to a loss of regulation in patterning and abundance of each cone type.

Our studies contribute to the understanding of cone development. We now know that along with *trβ2’s* role in the structural development of the cone layer, there are in fact alterations that extend to cone physiology and zebrafish behavior reflecting what was seen in immunohistochemistry. Future studies will examine *6BP*+*1* retinal circuitry to investigate downstream changes such as bipolar cell output and horizontal cell regulation. The *3BP* mutant lends itself to investigations on its lethal characteristics and what other pathways could be involved in this process.

## Materials and Methods

### Zebrafish

Zebrafish (*Danio rerio*) were kept in Aquatic Habitats benchtop systems (Pentair Aquatic Eco-Systems) and all procedures for breeding and experimental use were approved by the National Institute of Neurological Disorders and Stroke/National Institute on Deafness and Other Communication Disorders/National Center for Complementary and Integrative Health IACUC (ASP 1307, ASP 1227). Larvae between ages 5-7 days post fertilization (dpf) were kept in an incubator at 28° C in 3.5-inch Petri dishes filled with larval medium. The larval medium was made up of 60 mg/liter sea salt and 75 *μ*l/liter 1.5% methylene blue (Sigma-Aldrich Cat. No. 03978). At 8 dpf, the larvae were transferred to system nursery tanks (520-650 microOhms water, 28° C, pH 7.5-7.7) and fed Larval AP100 (Pentair Aquatic Eco-Systems) and live rotifers (*Brachionus Plicatilis*, Reed Mariculture) until experimental use at 12 dpf. Adults were kept in the same system environment as the 12 dpf larvae but were fed ground tetramin flakes and live rotifers.

### CRISPR/Cas9 and Phenotyping

Germ-line mutations of *trβ2* were generated by Cas9 mRNA and tr*β*2-gRNA co-injections into the one-cell stage eggs of *Tg(thrb2:tdTomato)q22* or *Tg(thrb:MA-YFP)q23* according to the method described previously [32]. The targeting sequence is 5’-GGCAACACAGCCAACCCTAT-3’ which resides in the first exon of *trβ2*, the only exon not shared by *trβ1*. The injected eggs were raised in 300 µM Phenylthiourea (PTU, Sigma-Aldrich, catalogue P7629) to blocks melanin synthesis between 12 hpf and 4dpf. At 4dpf, larvae with weak mosaic fluorescent expression of tdTomato or YFP in the photoreceptor layer were screened as potential carriers of germ-line mutations of *trβ2* and raised. The carriers of germ-line mutation of *trb2* were identified by in-crossing the potential carriers. The rationale was that *trβ2* mutants might have a fluorescence phenotype, as *trβ2* is a self-activating gene. Suspected mutant lines were in-crossed for multiple generations. To sort the fluorescence phenotypes, larvae were raised in 300 µM Phenylthiourea (PTU, Sigma-Aldrich, catalogue P7629). At the age of 4 dpf larvae were anesthetized in a 3.5-inch Petri dish with ∼0.5 ml of Tris buffered 0.4% tricaine in 45 ml of egg water before sorting on the basis of fluorescence in the eye’s photoreceptor layer

### Mosaic analysis of cone photoreceptor spatial arrangement

Chimera fish that were mosaic of *trβ2*+*/*+, *trβ2*+/-, and *trβ2-/-* cells were produced by injecting *trβ2* gRNA and cas9 mRNA into 1-cell stage embryos of a quadruple transgenic line, *Tg(gnat2:H2ACFP, sws1:H2AYFP, sws2:GFP, trβ2:tdTomato)*. Retinas were fixed at 28 dpf. 3D stack images were acquired using a confocal microscope that was equipped with 440, 515 and 561 nm laser for exciting CFP, YFP and tdTomato, respectively. Image analysis was conducted using Amira. Green cone nuclei were determined by subtracting red-, blue-, and UV-cone signals from the *gnat2*:H2ACFP fluorescence.

### Genotyping

To determine genotyping of the fish we performed sequencing from fish genomic DNA. The fish genomic DNA was extracted from whole larva or adult fish tail fin snip by KAPA express extract kit and PCR was performed using primers that span the target site of interest, the forward primer 5’-CATGGTGTAAGTGGCGGATATG-3’ and the reverse primer 5’-TCCACTGCATCTGAGAGAAATCC-3’. The PCR primer pairs were designed by the primer3 program (http://primer3.sourceforge.Net). PCR reactions were completed in 10 µl volumes containing 40 ng of genomic DNA, 1 µl of the forward and reverse primers at 10 µM, 1 µl of 10XPCR Buffer(100 mM Tris HC1(pH8.4)), 2.5 mM MgCl_2_, 2.5 mM dNTP mix and 0.2 U Taq DNA Polymerase. The thermal cycling conditions were as following: 94°C for 3 min, 35 cycles of 94°C for 30 sec, annealing temperature at 58°C for 30 sec and 72°C for 40 sec, and a final extension at 72°C for 3 min. PCR products were purified by using the AMPure XP system (Beckman Coulter, Biomek NX). Genomic PCR products were sequenced, the PCR primers were used for bidirectional sequencing using Big Dye Terminator Ready reaction mix according to manufacturer instructions (Applied Biosystems). Sequencing products were purified using the Agencourt CleanSEQ system on a Beckman Coulter, Biomek NX. Sequencing was performed on an ABI PRISM 3130 Automated sequencer (Applied Biosystems) and the sequencing results were analyzed using Mutation Surveyor v3.30 (Soft Genetics Inc., State College PA).

### Isolation and Perfusion of Eyes

Larvae at ages 5, 6, 7, and 12 dpf were isolated on a glass lantern slide, then transferred to a piece of nitrocellulose filter paper (Millipore, 0.45µm pore, Cat. No. HABP02500). With a 37 mm insect pin (Carolina Biological Supply), cuts were made behind the eyes to decapitate the larvae then longitudinally to make a dorsal-ventral cut between the eyes resulting in an isolated eye facing upward. Adults were decapitated with a fresh single-edged razor then longitudinally hemisected between the eyes. Attached tissues were removed from around the eye and the eye was placed upright on a piece of nitrocellulose filter paper. Under a dissecting scope, the cornea was sliced open to remove the lens and open the eyecup. In the recording chamber, the isolated eye was placed in an inverted lid of a 35-mm culture dish (ThermoFisher Scientific), with a disk of 41 µm nylon net filter (Millipore) covering the bottom to wick away perfusate. Larval eyes were perfused at 0.07 ml/min with minimal essential medium (MEM, Thermo FisherScientific, Cat. No. 11090-099, equilibrated with 95% O2, 5% CO2) using a syringe pump (New Era 500L, Braintree Scientific) and a 28-guage microfil syringe needle as an applicator (World Precision Instruments, MF28G67). The microfil applicator was positioned on the nylon mesh. Patch electrodes (3 µm tip) made using a Flaming/Brown microelectrode puller (Model P-87, Sutter Instrument, Novato, CA) and filled with 500 mM KCl were inserted transcorneally to record the massed cone PIII ERG signals. Adult eyecups were perfused with MEM (as above) at 0.3 ml/min. The perfusion applicator was placed directly in the eyecup to ensure retinal oxygenation. Shaved down microelectrodes (300 µm tips), placed in the eyecup, recorded cone-PIII signals (Nelson and Singla 2009). To record cone PIII ERG signals, L-Aspartate (Sigma-Aldrich, catalogue 11195, 20 mM larvae, 10mM adults) was added to MEM perfusate to block post-synaptic, glutamatergic, photoreceptor mechanisms.

### Electroretinogram Physiology

As in Nelson 2019 [16], light stimuli were obtained from a 150W OFR Xenon arc, shutter (Vincent Associates, Cat. No. LS6ZM2, 300 ms steps at 2.5-6.0 sec intervals), interference filters (330-650 nm, 40 nm increments, 20 nm half-width, Chroma Technology), metallic neutral density filters (7.5 log units, 0.5 log unit steps, Andover Corporation), computer-driven filter wheels, and liquid light guide (Sutter instruments, Lambda 10-3). Stimuli entered the epifluorescence port of an Olympus BX51 upright microscope (Olympus – Life Science Solutions) and were projected onto the retinas through UV-compliant objective lenses. A 10x UPlanFLN/0.3 projected stimuli on to isolated larval eyes and a 4x UPlanSApo/0.16 projected stimuli onto adult eyecups. A background beam infrared (IR, RG780 filter) was projected onto the eyes through a second light path as the ‘neutral’ background for infrared visualization of eye and electrode placement using an infrared camera (QImaging, Retiga-2000RV) and Metamorph (Molecular Devices). The microscope was positioned over the chamber with a translation stage (Sutter Instrument, MT-800) and microelectrodes were inserted into eyes (or eyecups) with a micro-positioner (Sutter Instrument, MPC-385). Microelectrode signals were amplified by 10,000 (World Precision Instruments, DAM80, 0.1 Hz-1k Hz bandpass), and digitized (2000 Hz) with an Axon instruments 1440A (Molecular Devices) using Clampex 10 software. The 280 ERGs within a spectral dataset were saved as a single Clampex file by using the averaging option and retaining all the elements of the average.

### Spectral Stimulus Protocol

The 17-min spectral protocol was a fixed sequence of 280, 300 msec, monochromatic light flashes of different irradiances delivered through house software. The 64 unique stimuli and 6 replicates created the spectral dataset, defined in Nelson 2019 [16] as a set of 70, 4X-averaged, ERG responses. Wavelengths were given in the order 650, 570, 490, 410, 330, 650, 610, 530, 450, 370 nm with 650 nm repeated as an index of response stability. At each wavelength 7 irradiances were delivered in 0.5 log unit increments, with brightness levels pre-adjusted to cover the anticipated response range from threshold to saturation. The interval between stimuli varied between 2.5 and 6 s, with the longer intervals separating the brighter stimuli. The protocol setting for maximal irradiances in log(quanta·µm^-2^·s^-1^) at each wavelength were 7.2 (650 nm), 6.3 (610 nm), 6.4 (570 nm), 6.3 (530 nm), 6.4 (490 nm), 6.1 (450 nm), 5.7 (410 nm), 5.7 (370 nm), and 5.2 (330 nm).

### Analysis of Spectral Data

Clampex files of spectral datasets were imported into Origin (various versions, Originlabs) for data analysis using scripts written in Origin Labtalk, where 4X replicates of the stimulus were averaged, noise was filtered (33 point, 16.5 msec running average), trough to peak amplitudes of PIII responses were calculated, and responses were associated with the wavelength and irradiance of stimulation. Datasets with unstable responses were over the recording period were not included.

The contributions of signals from different spectral types of cone to the PIII ERG response were determined using the method of Nelson et al [16]. Datasets from many eyes were normalized to the peak response within each and then combined into large ‘treatment-level’ datasets for 5-7 dpf larvae or for adults. These treatment-level datasets included thousands of amplitude-wavelength-irradiance points. Treatment-level datasets were fit to a spectral model (Eq.1) which extracted the physiological properties of the 4 spectral types of zebrafish cone. This provides a method to determine cone signals were altered by mutation, and in what way. The cone properties extracted were *Vmax*, the maximal amplitude contribution of the cone type to the PIII ERG, *k*, the irradiance required to half-saturate that cone’s response amplitude, and *λmax*, the spectral sensitivity maximum of that cone. The spectral model appears in equation 1.

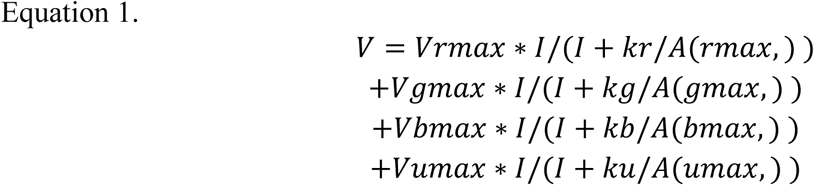

*V* is the cone PIII trough to peak signal amplitude, *I* is the quantal stimulus irradiance, and *λ* is the stimulus wavelength. The subscripts *r, g, b*, and *u* refer to red (*LWS* opsin expressing), green (*RH2* opsin expressing), blue (*SWS2* opsin expressing), and UV (*SWS1* opsin expressing) cones respectively. There are altogether 12 cone parameters that might be fit, however in many cases, for example datasets from mutants that lack signals from a cone type, the number that could be fit was less. This was evidenced by failure of the fitting algorithm (Levenburg Marquardt, as supplied by Originlabs) to find a solution within 100 iterations, or by the parameter encountering a boundary condition (Table 4). In this case a fixed value based on previous experience (Nelson and Singla, 2009) was substituted (Table 4).

**Table 4.** Boundaries and seed values for cone fit parameters.

| Fit parameter | Units | Fixed value | Lower bound | Upper bound |
| --- | --- | --- | --- | --- |
| $\log kr$ | $\text{quanta} \cdot \mu\text{m}^{-2} \cdot \text{s}^{-1}$ | 4.5 | 1 | 6 |
| $\log kg$ | $\text{quanta} \cdot \mu\text{m}^{-2} \cdot \text{s}^{-1}$ | 4.0 | 1 | 6 |
| $\log kb$ | $\text{quanta} \cdot \mu\text{m}^{-2} \cdot \text{s}^{-1}$ | 3.5 | 1 | 6 |

A(*λmax, λ*) is the Dartnall nomogram function (Dartnall) giving the spectral shape for zebrafish cone opsins. It is based in the observation that, on a reciprocal wavelength axis, the absorbance shapes of opsins with different absorbance maxima are closely similar. These nomogram shapes are expressed as order 8 polynomials. The polynomial of Hughes et al [33] is used for *r, g*, and *b* cones and the polynomial of Palacios et al [34] is used for *u* cones, which are slightly narrower in spectral shape. All nomograms are based in the spectral patch recordings of Palacios et al on *Giant Danio* [34], a species closely related to zebrafish. The polynomials are normalized, which has the effect of referring the fitted *k* (semisaturation irradiance) values for cones to *λmax*, the wavelength of maximal opsin absorbance. Nomogram coefficients are reproduced in Table 5.

**Table 5.** Polynomial coefficients for normalized cone opsin absorbance nomograms, A(λ_max_, λ).

| Coefficient | $r$, $g$, $b$ cones<br>Hughes et al, 1998<br>(Hughes *et al.*, 1998) | UV cones<br>Palacios et al, 1996<br>(Palacios *et al.*, 1996) |
| --- | --- | --- |
| $C_0$ | -93.262 | -48,139.554 |
| $C_1$ | 617.458 | 309,007.504 |
| $C_2$ | -2,815.735 | -770,875.542 |
| $C_3$ | 7,339.277 | 834,365.023 |
| $C_4$ | -10,714.545 | -57,657.482 |
| $C_5$ | 9,048.982 | -815,689.388 |
| $C_6$ | -4,410.667 | 871,189.984 |
| $C_7$ | 1,154.235 | -389,189.665 |
| $C_8$ | -125.738 | 66,989.122 |

### Confocal Imaging

Larval zebrafish are moved from egg water with methylene blue into a petri dish with 20mL of 300µM PTU at 1 day post fertilization. Larvae are sorted fluorescent and non-fluorescent phenotypes using the same methods as non-PTU larvae. At 5 days past fertilization larvae are placed into a separate petri dish and anesthetized with tricaine, as above. Low melting 0.8% agarose gel (Sigma-Aldrich, A0701) is melted on a hot plate to 35° C and a drop is placed into a Lab-Tek II 2-chambered cover-glass bottom well for microscopy (ThermoFisher Scientific, 155279PK). Each of the two chambers is divided in 4 by an insert, giving a total of 8 chambers. A larva is dropped into the well followed by a second drop of agarose. The larval eye is pressed flat against the bottom of the well with forceps and the agarose is allowed to cool and harden. A drop of tricaine was added to the top to keep the larvae anesthetized during imaging. A ZEISS LSM T-PMT confocal microscope was used with Zen image software (version 2.3). DPSS 561-10 (561 nm) laser was used to fluoresce tdTomato transgene. The mYFP transgene was imaged with an Argon 488 laser. A Z-stack image sets were collected at 25x and later analyzed using ImageJ (FIJI).

### Optomotor Response

Larvae between the ages of 5and 12 dpf were tested for optomotor response (OMR) in a 3.5-inch Petri dish placed directly on a laptop computer monitor (∼50 larvae per dish). The stimulus was generated using PsychoPy Python library package through PyCharm (v. 2018). Colored gratings swept from left to right on the monitor with a spatial frequency of 0.36 cycles/cm, size of 50, band width of 20mm, and phase of 0.03 (50mm/sec). Two stimuli were given: black/white and red/black. The black/white stimulus was given first, to isolate responsive larvae. The red/black stimulus was next presented to test for red-blind mutants.The stimulus lasted for 60 seconds and larvae were given a minute break between stimuli in a dark incubator. Suspected mutants were repeatedly tested. While the wavelength emission peaks for monitor colors are not specifically known, in general red is ∼600 nm; green, ∼550 nm, and blue, ∼450 nm. The monitors do not stimulate zebrafish UV cones. Videos and images were taken with an iPhone 8 Plus camera. Before and after images were analyzed using Mac Preview.

### Optokinetic Response

Larvae between the ages of 7 and 9dpf were tested for optokinetic response (OKR). A larva was placed in a drop of 4% methyl cellulose [20] on a glass lantern slide on the computer monitor. Pinwheel stimuli were created using the PsychoPy Python package through PyCharm (v. 2018). The pinwheel stimulus was made up of 12 alternating color wedges with the larvae at the center of the rotating stimulus. The colors included: black/white, red/blue, red/black, and green/black. The pinwheel rotated at 7.5 revolutions per minute. Correct eye movements were counted as a sweeping movement clockwise and snapping back counterclockwise. The numbers of correct eye movements were counted over a 60 second period.

### Immunohistochemistry of Larvae

Larvae fixed in 4% paraformaldehyde / 0.1M phosphate buffered saline (Electron Microscopy Sciences, catalogue 15710; Omnipur, catalogue 6505; PFA/PBS) for 30 min were dissected in a Petri dish filled with PBS using two 30-G disposable syringe needles to isolate retinas. The retinas were then transferred into 1.5 mL Eppendorf tubes (ThermoScientific, catalogue 3451) for subsequent blocking with lamb serum blocking buffer (Life Technolgies/Invitrogen, catalogue 6640), washing with Tween wash buffer (Invitrogen, catalogue 00-3005), and incubated with 1D4 anti zebrafish red-cone opsin primary antibody (AbCam, catalog ab5417, 1:100) and Alexa Fluor488 anti-mouse secondary antibody (Life Technologies/Invitrogen, catalogue 16070096, 1:100). The primary antibody was left for an incubation period of 3-4 days while the secondary was left for 1-2 days. Retinas were then mounted with Vectashield Antifade Mounting Medium with DAPI (Vector Laborotories, catalogue H-1200) between two coverslips separated by a spacer-square of black duct tape with a paper-punch hole in the middle to accommodate the retina.. The cover glass sandwich was pasted to a microscope slide using clear nail polish. The retinas were imaged on the same confocal microscope used for live imaging with the Argon (458, 488, 514), Diode (405-30) and DPSS 561-10 lasers. Images were analyzed with ImageJ (FIJI).

Adult eye cups were dissected and fixed in PFA/PBS for one hour and washed with PBS. Retinas were removed from the eye cups using No. 5 forceps to peel away the outside layers of the eye cup. Retinas were then placed in a 1.5 mL Eppendorfs for the same process of staining as with the larvae. A cocktail of zpr1 (ZFIN ID: ZDB-ATB-081002-43, 1:100) and rabbit anti-UV opsin primaries (1:100 dilution) were used in the first round of staining along with their respective secondaries Alexa Fluor594 anti-mouse (Thermo Fisher Scientific, catalog A-21135, 1:100) and Alexa Fluor405 anti-rabbit (Thermo Fisher Scientific, catalogue A-31556, 1:100) following the same timeline as previously mentioned. The anti UV opsin antibody was generously supplied by the Jeremy Nathans Laboratory, Johns Hopkins University) [35]. The second round of staining included 1D4 anti-red-opsin primary and Alexa Fluor488 anti-mouse secondary. Retinas were mounted on a coverglass fixed into place with a drop of 0. 8% agarose as above and mounted with VECTASHIELD Vibrance Antifade Mounting Medium. The samples were imaged and analyzed using the same methods as the larvae.

### Randomization/Blind, Inclusion/Exclusion, Sample Size Estimation

Heterozygous in-cress or heterozygous mutant spawns included 2-3 genotypes including the mutants, heterozygotes, WTs. Genotype was unknown when the fish were selected for physiological experimentation. Following physiological data collection, tail snip samples were given letter number names and sequenced without knowledge of physiological results. Optokinetic experiments were conducted without knowing the genotype of the larvae, but genotypes were known for the optomotor experiments. Live confocal imaging was conducted without known genotypes as we could not discern WTs from heterozygotes in the fluorescent group or any of the non-fluorescing fish. Genotypes of larval zebrafish were unknown before antibody staining, while adult zebrafish genotypes were known before staining.

We excluded physiological datasets with unstable response amplitudes over the course of the 17-minute protocol. Datasets that maintained stable response amplitudes were included. Unstable datasets could include either loss of electrode penetration or gradual declines/increases in amplitude. We excluded parameter fits that did not converge in the model and typical fixed values were substituted.

Previous studies have shown that 10 eyes/1000 amplitudes (spectral responses/data points) are enough to make the physiological distinctions expected [16].

## Acknowledgements

We would like to thank Jeremy Nathans for the UV opsin antibody and Douglas Forrest for his helpful discussions and references. The research was supported by the Intramural Program of the National Institute of Neurological Disorders and Stroke, National Institutes of Health, and by National Institutes of Health Grant EY14356 (to Rachel O. L. Wong)

## References

1. Ng, Lily; Hurley, James B; Dierks, Blair; Srinivas, Maya; Salto, Carmen; Vennstrom, Bjorn; Reh, Thomas A; Forrest Douglas. (2001). “A thyroid hormone receptor that is required for the development of green cone photoreceptors.” Nature genetics 27(1):94.

2. Liu, Yi-Wen; Lo, Li-Jan; Chan, Woon-Khiong. (2000). “Temporal expression and T3 induction of thyroid hormone receptors alpha1 and beta1 during early embryonic and larval development in zebrafish, Danio rerio.” Molecular and Cellular Endocrinology 159: 187–195.

3. Campi, Irene; Cammarata, Gabriella; Marzoli, Stefania B; Beck-Peccoz, Paolo; Santarsiero, Diletta; Dazzi, Davide; Bottari de Castello, Alessandra; Taroni, Elena G; Viola, Francesco; Mian, Caterina; Watutantrige-Fernando, Sara; Pelusi, Carla; Muzza, Marina; Maffini, Maria A; Persani, Luca. (2017). “Retinal photoreceptor functions are compromised in patients with resistance to thyroid hormone syndrome (RTHβ).” The Journal of Clinical Endocrinology & Metabolism 102(7):2620–2627.

4. Weiss, Avery H; Kelly, John P; Bisset, Darren; Deeb, Samir S. (2012). “Reduced L-and M-and increased S-cone functions in an infant with thyroid hormone resistance due to mutations in the THRβ2 gene.” Ophthalmic genetics 33(4):187–195.

5. Sjöberg, M. and B. Vennström (1995). “Ligand-dependent and-independent transactivation by thyroid hormone receptor beta 2 is determined by the structure of the hormone response element.” Molecular and Cellular Biology 15(9):4718–4726.

6. Ng, Lily; Lu, Ailing; Swaroop, Alok; Sharlin, David S; Swaroop, Anand; Forrest, Douglas. (2011). “Two transcription factors can direct three photoreceptor outcomes from rod precursor cells in mouse retinal development.” Journal of Neuroscience 31(31):11118–11125.

7. Darras, Veerle M; Van Herck, Stijn LJ; Heijlen, Marjolein; De Groef, Bert. (2011). “Thyroid hormone receptors in two model species for vertebrate embryonic development: chicken and zebrafish.” Journal of Thyroid Research 2011.

8. Deeb, Samir S. (2006). “Genetics of variation in human color vision and the retinal cone mosaic.” Current opinion in genetics & development 16(3):301–307.

9. Marelli, Federica; Carra, Silvia; Cotelli, Franco; Peeters, Robin; Chatterjee, Krishna; Persani, Luca. (2016). “Patterns of thyroid hormone receptor expression in zebrafish and generation of a novel model of resistance to thyroid hormone action.” Molecular and Cellular Endocrinology 424: 102–117.

10. Suzuki, Sachihiro C; Bleckert, Adam; Williams, Philip R; Takechi, Masaki; Kawamura, Shoji; Wong, Rachel OL. (2013). “Cone photoreceptor types in zebrafish are generated by symmetric terminal divisions of dedicated precursors.” Proceedings of the National Academy of Sciences: 201303551.

11. Chinen, Akito; Hamaoka, Takanori; Yamada, Yokihiro; Kawamura, Shoji. (2003). “Gene duplication and spectral diversification of cone visual pigments of zebrafish.” Genetics 163(2):663–675.

12. Mitchell, Diana M; Stevens, Craig B; Frey, Ruth A; Hunter, Samuel S; Ashino, Ryuichi; Kawamura, Shoji; Stenkamp, Deborah L. (2015). “Retinoic acid signaling regulates M differential expression of the tandemly-duplicated long wavelength-sensitive cone opsin genes in zebrafish.” PLoS genetics 11(8):e1005483.

13. DuVal, M. G. and W. T. Allison (2018). “Photoreceptor Progenitors Depend Upon Coordination of gdf6a, thrβ, and tbx2b to Generate Precise Populations of Cone Photoreceptor Subtypes.” Investigative ophthalmology & visual science 59(15):6089–6101.

14. Jones, I., et al. (2003). “The thyroid hormone receptor β gene: structure and functions in the brain and sensory systems.” Thyroid 13(11):1057–1068.

15. Allison, W Ted; Barthel, Linda K; Skebo, Kristina M; Takechi, Masaki; Kawamura, Shoji; Raymond, Pamela A. (2010). “Ontogeny of cone photoreceptor mosaics in zebrafish.” Journal of Comparative Neurology 518(20):4182–4195.

16. Nelson, Ralph F; Balraj, Annika; Suresh, Tara; Torvund, Meaghan; Patterson, Sara S. (2019) “Strain variations in opsin peaks in situ during zebrafish development.” Visual neuroscience 36, E010 DOI: https://doi.org/10.1017/S0952523819000075.

17. D’Agati, Gianluca; Beltre, Rosanna; Sessa, Anna; Burger, Alexa; Zhou, Yi; Mosimann, Christian; White, Richard M. (2017). “A defect in the mitochondrial protein Mpv17 underlies the transparent casper zebrafish.” Developmental Biology 430(1):11–17.

18. Orger, M. B. and H. Baier (2005). “Channeling of red and green cone inputs to the zebrafish optomotor response.” Visual neuroscience 22(03):275–281.

19. Krauss, A. and C. Neumeyer (2003). “Wavelength dependence of the optomotor response in zebrafish (Danio rerio).” Vision research 43(11):1275–1284.

20. Brockerhoff, Susan E; Hurley, James B; Janssen-Bienhold, Ulrike; Neuhauss, SC; Driever, Wolfgang; Dowling, John E. (1995). “A behavioral screen for isolating zebrafish mutants with visual system defects.” Proceedings of the National Academy of Sciences 92(23):10545–10549.

21. Shibusawa, Nobuyuki; Hollenberg, Anthony N; Wondisford, Fredric E. (2003). “Thyroid hormone receptor DNA binding is required for both positive and negative gene regulation.” Journal of Biological Chemistry 278(2):732–738.

22. Tsujimura, Taro; Hosoya, Tomohiro; Kawamura, Shoji. (2010). “A single enhancer regulating the differential expression of duplicated red-sensitive opsin genes in zebrafish.” PLoS genetics 6(12):e1001245.

23. Takechi, M. and S. Kawamura (2005). “Temporal and spatial changes in the expression pattern of multiple red and green subtype opsin genes during zebrafish development.” Journal of Experimental Biology 208(7):1337–1345.

24. Mackin, Robert D; Frey, Ruth A; Gutierrez, Carmina; Farre, Ashley A; Kawamura, Shoji; Mitchell, Diana M; Stenkamp, Deborah L. (2019). “Endocrine regulation of multichromatic color vision.” Proceedings of the National Academy of Sciences 116(34):16882–16891.

25. Harvey CB, Williams GR. (2002). “Mechanism of thyroid hormone action.” Thyroid. 2002;12:441–446.

26. Baier, Herwig. (2000). “Zebrafish on the move: towards a behavior–genetic analysis of vertebrate vision.” Current opinion in neurobiology 10(4):451–455.

27. Woods, Andrew; Lun Yuen, Ka; S. Karvinen, Kai. (2007). Characterizing crosstalk in anaglyphic stereoscopic images on LCD monitors and plasma displays. Journal of the Society for Information Display. 15. 10.1889/1.2812989.

28. M NN Srinivas, Maya; Ng, Lily; Liu, Hong; Jia, Li; Forrest, Douglas. (2006). “Activation of the blue opsin gene in cone photoreceptor development by retinoid-related orphan receptor β.” Molecular endocrinology 20(8):1728–1741.

29. Li, Yong N; Matsui, Jonathan I; Dowling, John E. (2009). “Specificity of the horizontal cell-photoreceptor connections in the zebrafish (Danio rerio) retina.” Journal of Comparative Neurology 516(5):442–453.

30. Renninger, Sabine L, Gesemann, Matthias, Neuhauss, Stephan CF (2011). “Cone arrestin confers cone vision of high temporal resolution in zebrafish larvae.” European Journal of Neuroscience 33(4):658–667.

31. Raymond, Pamela. A, Barthel, Linda K (2004). “A moving wave patterns the cone photoreceptor mosaic array in the zebrafish retina.” International Journal of Developmental Biology 48(8-9): 935–945.

32. Jao, Li-En, Wente, Susan R, and Chen, Wenbiao (2013). “Efficient multiplex biallelic zebrafish genome editing using a CRISPR nuclease system.” Proceedings of the National Academy of Sciences 110(34):13904–13909.

33. Hughes, Alan., Saszik, Shannon, Bilotta, Joseph, Demarco, Paul J Jr., Patterson, Warren F 2nd (1998). “Cone contributions to the photopic spectral sensitivity of the zebrafish ERG.” Visual Neuroscience 15(6):1029–1037.

34. Palacios, Adrian G., Goldsmith, Timothy H, Bernard, Gary D (1996). “Sensitivity of cones from a cyprinid fish (Danio aequipinnatus) to ultraviolet and visible light.” Visual Neuroscience 13(3):411–421.

35. Luo, Wenqin, Williams, John, Smallwood, Philip M, Touchman, Jeffrey W, Roman, Laura M, Nathans, Jeremey (2004). “Proximal and distal sequences control UV cone pigment gene expression in transgenic zebrafish.” J Biol Chem 279(18):19286–19293.

